# Creating Standards for Evaluating Tumour Subclonal Reconstruction

**DOI:** 10.1101/310425

**Authors:** Adriana Salcedo, Maxime Tarabichi, Shadrielle Melijah G. Espiritu, Amit G. Deshwar, Matei David, Nathan M. Wilson, Stefan Dentro, Jeff A. Wintersinger, Lydia Y. Liu, Minjeong Ko, Srinivasan Sivanandan, Hongjiu Zhang, Kaiyi Zhu, Tai-Hsien Ou Yang, John M. Chilton, Alex Buchanan, Christopher M. Lalansingh, Christine P’ng, Catalina V. Anghel, Imaad Umar, Bryan Lo, William Zou, DREAM SMC-Het Participants, Jared T. Simpson, Joshua M. Stuart, Dimitris Anastassiou, Yuanfang Guan, Adam D. Ewing, Kyle Ellrott, David C. Wedge, Quaid D. Morris, Peter Van Loo, Paul C. Boutros

## Abstract

Tumours evolve through time and space. Computational techniques have been developed to infer their evolutionary dynamics from DNA sequencing data. A growing number of studies have used these approaches to link molecular cancer evolution to clinical progression and response to therapy. There has not yet been a systematic evaluation of methods for reconstructing tumour subclonality, in part due to the underlying mathematical and biological complexity and to difficulties in creating gold-standards. To fill this gap, we systematically elucidated the key algorithmic problems in subclonal reconstruction and developed mathematically valid quantitative metrics for evaluating them. We then created approaches to simulate realistic tumour genomes, harbouring all known mutation types and processes both clonally and subclonally. We then simulated 580 tumour genomes for reconstruction, varying tumour read-depth and benchmarking somatic variant detection and subclonal reconstruction strategies. The inference of tumour phylogenies is rapidly becoming standard practice in cancer genome analysis; this study creates a baseline for its evaluation.

---

Most tumours arise from a single ancestral cell, whose genome acquires one or more somatic driver mutations^1,2^, which give it a fitness advantage over its neighbours by manifesting hallmark characteristics of cancers^3^. This ancestral cell and its descendants proliferate, ultimately giving rise to all cancerous cells within the tumour. Over time, they accumulate mutations, some leading to further fitness advantages. Eventually local clonal expansions can create subpopulations of tumour cells sharing subsets of mutations, termed *subclones.* As the tumour extends spatially beyond its initial location, spatial variability can arise as different regions harbour independently-evolving tumour cells with distinctive genetic and non-genetic characteristics^4–9^.

DNA sequencing of tumours allows quantification of the frequency of specific mutations based on measurements of the fraction of mutant sequencing reads, the copy number state of the locus and the tumour purity^10,11^. By aggregating these noisy frequency measurements across mutations, a tumour sample’s subclonal architecture can be reconstructed from bulk sequencing data^6,11^. Subclonal reconstruction methods have proliferated rapidly in recent years^12–15^, and have revealed key characteristics of tumour evolution^4,7,16–20^, spread^21–23^ and response to therapy^24,25^. Nevertheless, there has been no rigorous benchmarking of the relative or absolute accuracy of approaches for subclonal reconstruction.

There are several reasons why such benchmarking has not yet been performed. First, it is difficult to identify a gold-standard truth for subclonal reconstruction. While single-cell sequencing could provide ground truth, it has pervasive errors^26^, and existing DNA-based datasets do not have sufficient depth and breadth to adequately assess subclonal reconstruction methods. Alternatively, gold-standard datasets may be generated using simulations, but existing tumour simulation methods like BAMSurgeon^27^ neither create representative subclonal populations nor phase simulated variants, which can be exploited in subclonal reconstruction^6,10^. Second, it is unclear how subclonal reconstruction methods should be scored, even in the presence of a suitable gold-standard. For example, one key goal of reconstruction is identification of the mutations present in each subclonal lineage. Metrics are needed that penalise errors both in the number of subclonal lineages and in the placement of mutations across them. Third, subclonal reconstruction methods have only been developed in recent years; few groups have equal expertise with multiple tools. Algorithm developers themselves are typically experts in parameterizing their own algorithms; an unbiased third-party is needed compare different methods, each run with expert parameterization.

To fill this gap, we developed a crowd-sourced benchmarking Challenge: The ICGC-TCGA DREAM Somatic Mutation Calling Tumour Heterogeneity Challenge (SMC-Het). Challenge organisers simulated realistic tumours, developed robust scoring metrics and created a computational framework to facilitate unbiased method evaluation. Challenge participants then created re-distributable software containers representing their methods. These containers were run by the Challenge organizers in an automated pipeline on a series of test tumours never seen by the Challenge participants. Here, we describe the creation of quantitative metrics for scoring tumour subclonality reconstructions and of novel tools for simulating tumours with realistic subclonal architecture. Finally, we use these characterise the sensitivity of subclonal reconstruction methodologies to somatic mutation detection algorithms and technical artefacts.

## Results

### How should subclonal reconstruction methods be evaluated?

Subclonal reconstruction is a complex procedure that involves estimating many attributes of the tumour including its purity, number of lineages, lineage genotypes and the phylogenetic relationships amongst lineages. We structured our evaluation of these attributes into three categories (**Figure 1**). Sub-challenge 1 (SC1) quantify the ability of an algorithm to reconstruct global characteristics of tumour composition. Specifically, it evaluates each algorithm’s predictions of the total fraction of cells that are cancerous (tumour purity; SC1A), the number of subclonal lineages (SC1B) and for each subclone the fraction of cells (cellular prevalence) and number of mutations associated with it (SC1C). Sub-challenge 2 (SC2) evaluates how accurately each algorithm assigns individual single nucleotide variants (SNVs) to each subclonal lineage. It evaluates both their single-best guess at a hard assignment of SNVs to lineages (SC2A) and soft assignments represented through co-clustering probabilities *(i.e.* the probability that two SNVs are in the same lineage; SC2B). Finally, sub-challenge 3 (SC3) evaluates the ability of algorithms to recover the phylogenetic relationships between subclonal lineages, again both as a single hard assignment (SC3A) and as a soft assignment (SC3B). Taken together, these subchallenges define seven specific sub-challenges of SMC-Het, each corresponding to a specific outputs upon which subclonal reconstruction methods can be benchmarked (**Online Methods**).

**Figure 1.**
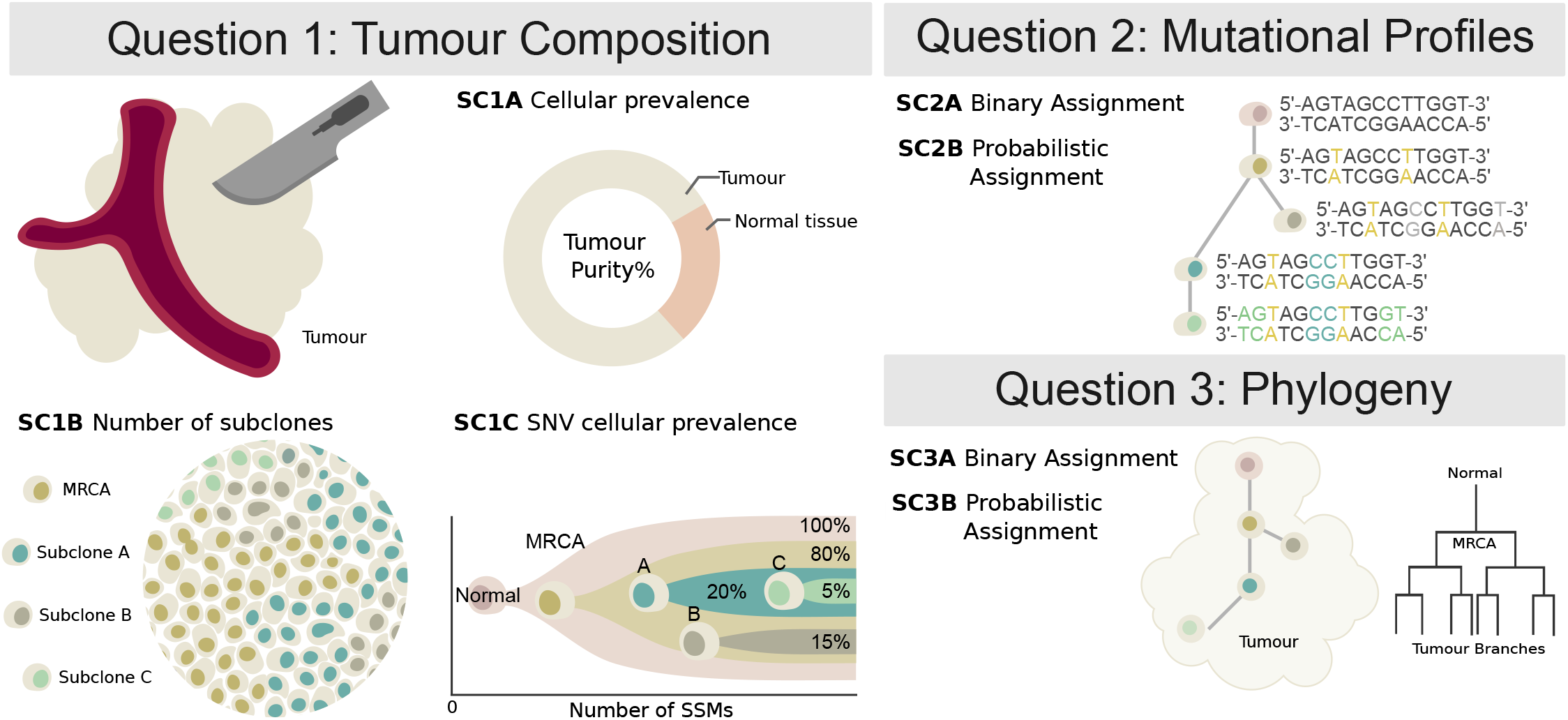
Features of tumour subclonal reconstruction. Overview of the key performance aspects of subclonal reconstruction algorithms, grouped into three broad areas covered by three key questions: (SC1) ‘What is the composition of the tumour?’ This involves quantifying its purity, the number of subclones, and their prevalence and mutation loads; (SC2) ‘What are the mutational characteristics of each subclone?’ This can be answered both with a point-estimate and a probability profile, *i.e.* a hard or probabilistic assignments of mutations to subclones, respectively; (SC3) ‘What is the evolutionary relationships amongst tumour subclones?’ This again can be answered with both a point-estimate and a probability profile. MRCA: most recent common ancestor.

To quantify the accuracy of these seven outputs, we considered several candidate scoring metrics, all bound between zero (very poor performance) and one (perfect performance). Appropriate metrics for SC1 were trivially identified (**Online methods**), but SC2 and SC3 required us to modify existing metrics and develop new ones. Specifically, because SC2B and SC3B are based on pairwise probabilities of coclustering, we were unable to use clustering quality metrics designed for hard clustering nor those that require explicit estimation of the number of clusters, such as normalised mutual information (also known as the V-measure^28^).

As SC2 and SC3 involve assigning mutations to subclonal lineages, we required candidate metrics to satisfy three conditions^28^:

1. The score decreases as the predicted number of subclonal lineages diverges from the true number of subclonal lineages.
2. The score decreases as the proportion of mutations assigned to incorrect subclonal lineages (predicted subclonal lineages that do not correspond to the true subclonal lineage) increases.
3. The score decreases as the proportion of mutations assigned to noise subclonal lineages (predicted subclonal lineages that do not correspond to any true subclonal lineage) increases.

Moreover, metrics for evaluating cluster assignments have a number of desirable properties^28^. We identified a set of these applicable to each task (**Online Methods**), used a simulation framework to assess how well a candidate metric satisfies them. We identified four complementary metrics that satisfy all three properties: Matthew’s Correlation Coefficient (MCC), Pearson’s Correlation Coefficient (PCC), area under the precision recall curve (AUPR) and average Jensen-Shannon divergence (AJSD; **Supplementary Figure 1**).

To further refine this set, we tested their behaviour relative to subclonal reconstruction errors related to parent *vs.* child and parent *vs.* cousin relationships, and splitting or merging of individual nodes (**Online methods**). Nine experts ranked the overall severity of up to eight error cases for each of 30 tree topologies, providing 2,088 total expert rankings. We then simulated each error case and scored it with all candidate metrics (**Figure 2a-d**). Importantly for SC3, we added one metric, the Clonal Fraction (CF), which scores the accuracy of the predicted fraction of mutations assigned to the clonal peak. Unlike SC2, which scores mutation assignment, *i.e.* genotyping of the (sub)clones, SC3 scores tree topology, which implies an ordering of events. The clonal fraction was designed to capture expert knowledge that emerged from the expert ranking: experts tended to favour the merging of two subclonal clusters over merging of the clonal cluster with a subclonal cluster, which was not captured by other metrics. The fraction of (sub)clonal mutations is indeed a biologically relevant measure that varies widely across cancer types^29^. Given that our metric rankings are based on subjective expert viewpoints, we have made our ranking system available online to allow others to create their own rankings and compare them to ours or use them to fine-tune scoring metrics for their own applications (https://mtarabichi.shinyapps.io/SMCHET).

**Figure 2.**
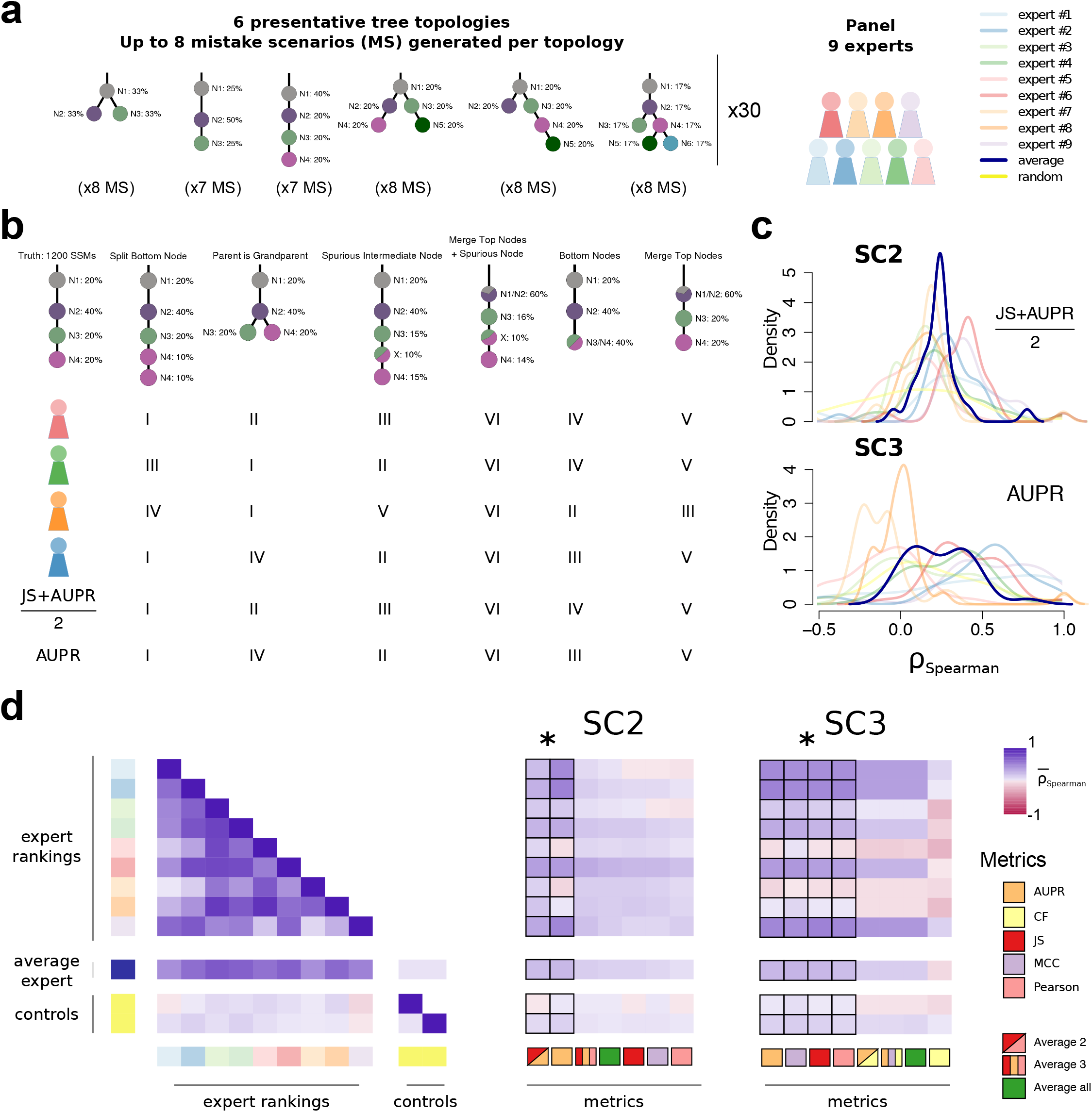
Quantifying performance of subclonal reconstruction algorithms. **(a) Tree topologies and mistake scenarios**. For each of 30 tree topologies with varying number of clusters and ancestral relationships, 7-8 mistake scenarios (MS) were derived and scored using the identified metrics for SC2 and SC3. For each tree topology a panel of 9 experts independently ranked the mistake scenarios from best to worse. **(b) Expert ranking**. One tree topology is shown with 6 of the 7 mistake scenarios together with the ranks of four experts and two of the metrics. The trivial allin-one case, i.e. identifying only one cluster is not shown and correctly ranked last by all metrics and experts. **(c) Density distributions of Spearman’s correlations between metrics and experts across tree topologies**. For SC2 and SC3, we show the Spearman’s correlations between JS+AUPR/2 and the experts, and AUPR and the experts, respectively. **(d) All average correlations between experts and metrics for SC2 and SC3**. Heatmaps of average Spearman’s correlations across tree topologies between experts and metrics for SC2 and SC3. Controls are randomised ranks. Asterisks show equivalent metrics (non-significantly better or worse according to a Mann-Whitney U test p>0.05 but better than the others p<0.01).

Between-expert agreement, measured as pairwise rank correlations (0.52 ± 0.22), were much higher than metrics-expert agreement (for SC2B, 0.14 ± 0.12; for SC3B, 0.12 ± 0.15; **Figure 2e**). Subsets of metrics were highly correlated (JS, Pearson and MCC; range: 0.97-0.99), whereas others were less correlated (AUPR, JS/Pearson/MCC and CF; range: 0.47-0.78). We reasoned that less correlated metrics might capture complementary aspects of the reconstructions and derived additional metrics combining the best of them, as well as an average of all (**Figure 2e**). For SC2, the average of two metrics 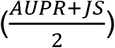 and AUPR was significantly better correlated to experts than any individual metric (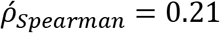; **Figure 2c,e**). For SC3, AUPR, MCC, Pearson and JS were comparable and significantly better than the other metrics (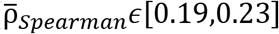. We chose Pearson for subsequent analysis as it allows for assessment with a non-binary truth. The resulting expert rankings and quantitative comparisons provide a basis for future development of improved scoring metrics.

### Simulating accurate subclonal tumour genomes

We elected to use simulated tumour data to run SMC-Het. The key reasons were the unavailability of deep single-cell DNA sequencing data as a gold-standard, the lack of single-cell sequencing data that match arbitrary tree structures and characteristics, the ability to simulate a large number of tumours at low-cost and the demonstrated ability of tumour simulations to recapitulate complex sequencing error profiles^27^. We elected to use the BAMSurgeon tool created for the earlier SMC-DNA Challenges^27,30^, which creates tumours with accurate SNVs, indels and small genomic rearrangements at varying allelic fractions. However, that version of BAMSurgeon lacked a number of key features for our purpose. We added five major features: (1) phasing of variants, (2) large-scale allele-specific copy number changes (including whole-genome duplications), (3) translocations, (4) trinucleotide SNV signatures and (5) replication-timing effects (**Figures 3, 4**). We describe each of these briefly.

**Figure 3.**
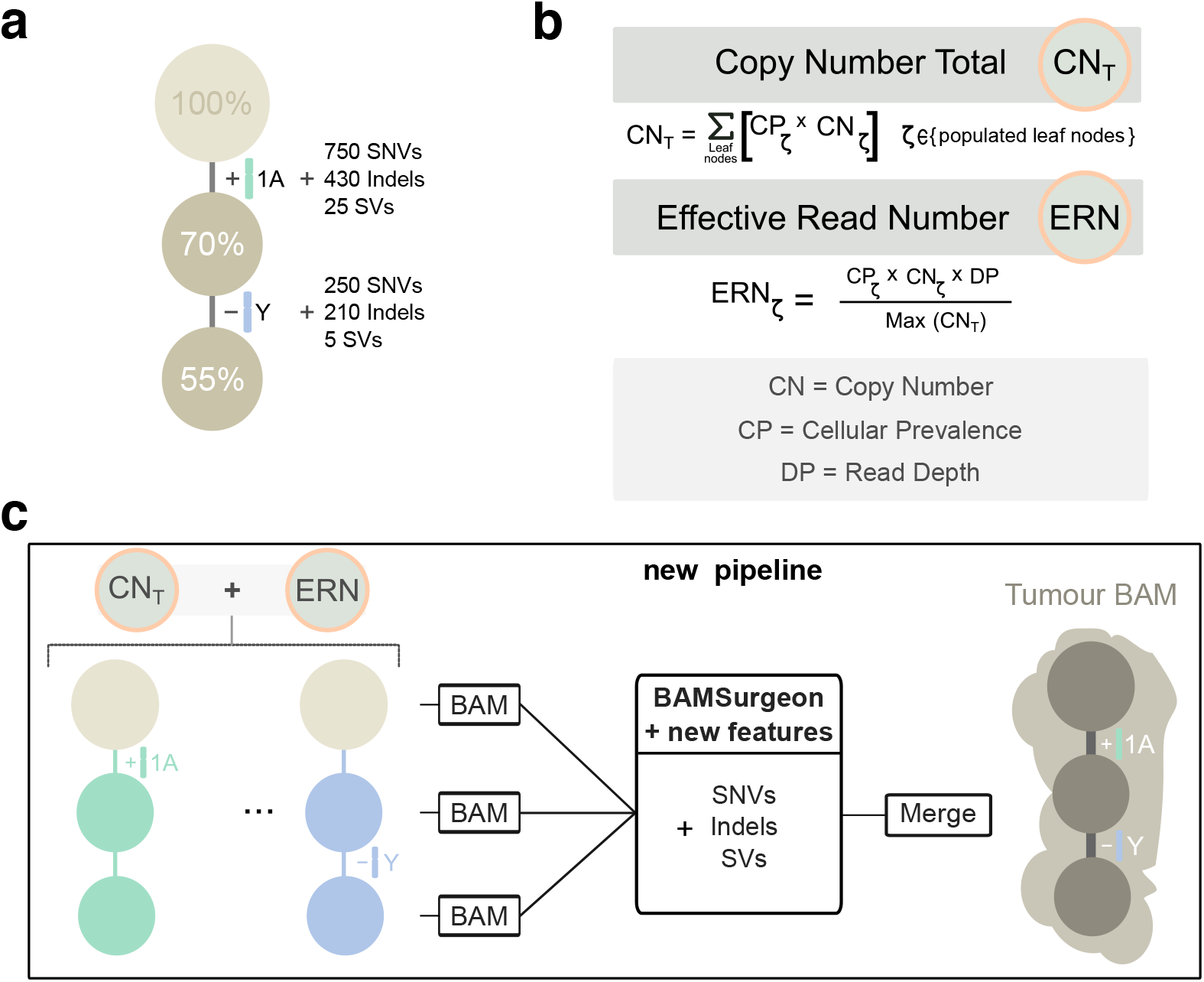
Simulating subclonal CNAs in Tumour BAM files and spiking somatic mutations. Example case of read number adjustment to simulate subclonal copy number aberrations (CNAs). **(a) Desired structure of the tumour being simulated. (b) Read number adjustment calculations**. The copy number total (CNT) for each chromosome is its copy number by adjusted by node cellular prevalence summed across all nodes. The maximum CNT across the genome is retained to normalise copy number for all chromosomes. The number of reads assigned to each chromosome at each node (the chromosome’s effective read number) is then computed as the product of the node’s cellular prevalence, the chromosome’s copy number, and the total tumour depth normalised by the maximum CNT. **(c) Separation per chromosome phase and per node and new pipeline to simulate tumour BAM files with underlying intra tumour heterogeneity**. The first tumour clone (70% CP) has a gain in one copy (referred to as copy A) of chromosome 1 and one of its descendant subclones (55% CP) bears a loss of the Y chromosome. After adjusting read number for CNAs in each BAM corresponding to a node, BAMSurgeon spikes in additional mutations including the new features (complex structural variants, SNVs with trinucleotide contexts and replication timing effects, *etc.*), and then merges the extracted reads into a final tumour BAM file.

**Figure 4.**
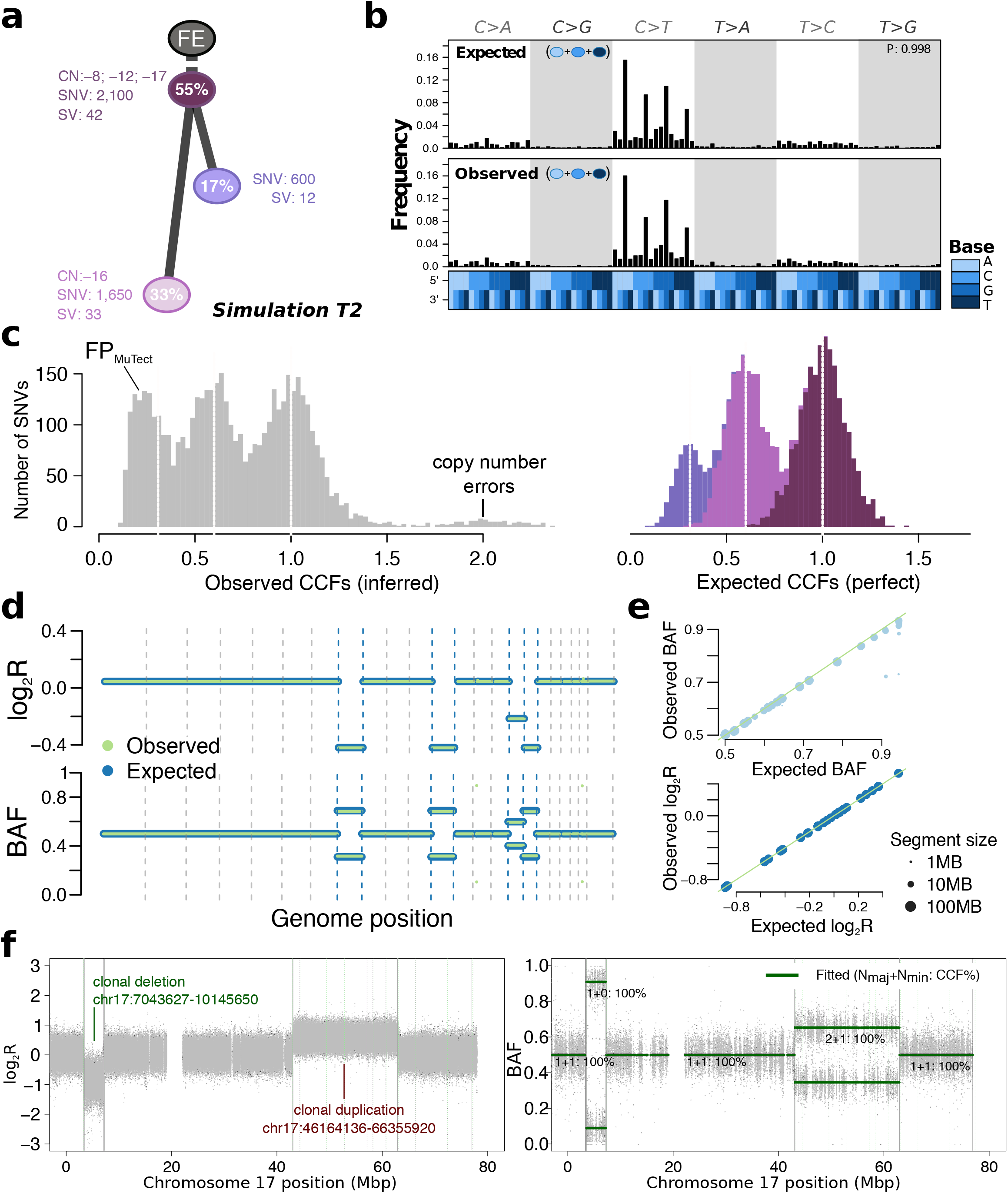
Simulated realistic tumour genomes. **(a) Tumour design**. Simulation T2 with 55% purity (fraction of cancer cells) and two subclones. Whole-chromosome copy number events (e.g. clonal loss of chromosomes 8, 12 and 17), number of SNVs and SVs are shown for each node. **(b) Single nucleotide variant trinucleotide contexts**. Observed vs. expected frequencies of trinucleotide contexts in the SNVs. **(c) Population frequency (cancer cell fraction, CCF) of the variants for T2**. Observed vs. expected CCF distributions; false positive SNVs due to mutation calling as well as copy-number errors lead to errors in the inferred CCFs. **(d) Observed (green) vs. expected (blue) logged coverage ratio (LogR) and B-allele frequencies (BAF) of copy number segments along the genome for T2 (e)** Observed vs. expected BAF and logR across all segments and across all simulations. **(f) Simulation of sub-chromosomal copy number events and rearrangements**. LogR and BAF tracks showing how one large deletion and one large duplication simulated on chromosome 17 are correctly being called. Structural variants as called by Manta (Online methods) are shown as vertical lines, true positives are at the breakpoints defining the copy number events.

#### Phasing of mutations

To correctly simulate a tumour, it is critical that genetic variants – both somatic and germline – are fully phased, as they are in real genomes. Without phasing, allele-specific copy number changes cannot be simulated correctly and will lead to incorrect B-allele frequencies and allele-specific copy number calls, amongst other errors. To achieve correct and complete phasing, we leveraged NGS data from a trio of individuals from the Genome-in-a-Bottle consortium (**Supplementary Figure 2a-e**) and created the PhaseTools package to accurately phase heterozygous variants identified in these data (**Online Methods**). The final result of this process is two BAM files per chromosome, each representing a single parental copy.

#### Simulation of a tumour BAM with underlying tree topology

(**Figure 3a**). To simulate a tumour BAM starting from the fully phased genome, we assigned subsets of the reads to each tree node, generating down-sampled BAM files. To simulate whole chromosome copy number events, we adjusted the proportion of reads assigned to each node of the tree (**Figure 3b**; see below). Then, BAMSurgeon was used on each sub-BAM to simulate mutations, including SNVs, indels and SVs (**Figure 3c**). This strategy allowed us to efficiently and reliably simulate copy number changes of arbitrary size and add specific mutations on each allelic copy. Finally, these sub-BAMs were merged to produce the final BAM. By contrast, when we used the subclonally-naive BAMSurgeon, copy number inference was incorrect (**Supplementary Figure 2f,g**). After adding subclonal mutations only by specifying the VAF (*i.e.* without phasing or subsampling BAM files) SNVs that occurred after duplications or deletions often appeared at the wrong frequency (**Supplementary Figure 2h**).

#### Whole arm and whole genome copy number changes

To allow changes in copy number of entire chromosomes and whole-genome ploidy changes *(e.g.* whole genome duplications, present in 30-50% of human cancers^31–33^), we developed a method to account for gains or losses of any chromosome, including sex chromosomes based on bookkeeping of reads assigned to each node. Given a tumour design structure (**Figure 3b**), reads from the phased genomes were further split into individual subpopulations (sub-BAMs for leaf nodes) that make up the tumour in proportion to the copy number state of the region they aligned to and the cellular prevalence of their node. The extracted and modified reads were merged to generate a final BAM file (**Figure 3c**).

#### Translocations and large-scale SVs

As the prior BAMSurgeon functionality could not reliably simulate SVs larger than 30 kbp or any translocations due to its use of assembly, we extended it to simulate translocations, inversions, deletions and duplications of arbitrary size. This required a new approach of creating a simulated translocation that accurately reflects the expected pattern of discordant read pair mappings and split reads (**Online Methods**). This also allows us to simulate translocations, which were not included in the SMC-DNA simulated data challenges^30^. The ability to simulate translocations combined with adjustments to read coverage makes the simulation of arbitrarily large and complex SVs possible.

#### Trinucleotide mutation profile and replication timing

Single nucleotide mutations are not uniformly distributed throughout cancer genomes. They are biased both regionally and locally^34^. Mutations result from specific mutagenic stresses, which can induce biased rates of occurrence at specific trinucleotide contexts^35^. Replication-timing bias refers to the increase in the mutation rate of regions of the genome that replicate late in the cell cycle^34^. To resolve this issue, we created an extensible approach as part of BAMSurgeon. Each nucleotide in the genome is weighted according to its trinucleotide context, replication timing and the set of mutational signatures. Bases are then sampled from the genome until the expected trinucleotide spectrum is reached (**Online Methods**). BAMSurgeon can handle arbitrary mutational signatures, replication timing data at any resolution and any arbitrary type of locational bias in mutational profiles.

#### Selection

Our framework for picking selecting point mutations can easily be extended to incorporate other biases in mutation frequency or location such as selection. Although explicit tumour growth models remain an area of active development^36–38^ and discussion^39,40^ we sought to illustrate this functionality using a recent model of 3D tumour growth that shows selection is reflected in VAF distributions across 3D tumour subvolumes^37^. We obtained VAFs from this simulator at five different levels of selection. For each level of selection, we simulated one 3D tumour and the resection of three tumour subregions. These were taken as basis for our simulator to generate 15 tumour BAM files in which the spiked-in SNVs and their VAF were directly derived from the tumour growth models. The VAFs of the genotyped SNVs allowed accurate inference of the selection input parameter (**Supplementary Figure 2i, Online methods**), while also incorporating tri-nucleotide signatures and replication timing effects. By contrast, we were unable to recover the signature of selection with MuTect SNV calls, suggesting that more than three tumour regions might be needed to detect selection through this method when significant variant detection errors are present, emphasizing the utility of simulated tumour BAMs in algorithm and model assessment (**Online methods**).

Each of the simulated features was verified by comparing simulated to designed values: observed to expected measurements in the BAMs (**Online Methods, Supplementary Figure 3**). Starting from a tumour design (**Figure 4a**) we systematically and quantitatively compared observed and expected trinucleotide context (**Figure 4b**), cancer cell fraction (**Figure 4c**) and copy number segment logR ratios and B-allele frequencies (**Figure 4d,e**). These were reviewed across all simulations to verify simulated data. These results also confirmed that BAMSurgeon can now generate complex sub-chromosomal events, including large deletions or duplications (**Figure 4f**).

### General features of subclonal reconstruction

We next sought to quantify how different factors impact subclonal reconstruction. We therefore simulated five tumours derived from different tissue types (prostate, lung, chronic lymphocytic leukaemia, breast and colon) from published subclonal structures (**Supplementary Figure 3**). We also analysed a real tumour (PD4120) sequenced at 188x coverage with a high-quality consensus subclonal reconstruction based on the full-depth tumour^41^ as the gold-standard.

For each of these six tumours, we then down-sampled each tumour sequence to create a titration series in raw read-depth of 8x, 16x, 32x, 64x and 128x coverage. For each of the 30 resulting tumour-depth combinations, we identified subclonal copy number aberrations (CNAs) using Battenberg^6^, both with down-sampled tumours and with tumours at the highest possible depth to assess the influence of CNA detection accuracy, yielding 60 tumour-depth-CNA combinations. For each of these combinations, we identified somatic SNVs using four algorithms (MuTect^42^, SomaticSniper^43^, Strelka^44^, and MutationSeq^45^), as well as the perfect somatic SNV calls for the simulated tumours, yielding 290 synthetic tumour-depth-CNA-SNV combinations. We also applied these pipelines to the real PD4120 BAM (except those involving of perfect SNV calls) resulting in 40 additional depth-CNA-SNV combinations based on a real tumour, for a total of 290 combinations. The somatic SNV detection algorithms were selected to span a range of variant calling approaches: SomaticSniper uses a Bayesian approach, MuTect and Strelka model allele frequencies while MutationSeq predicts somatic SNVs with an ensemble of four classifiers trained on a gold-standard dataset. Finally, subclonal reconstruction was then carried out on each of these using two algorithms (PhyloWGS^13^ and DPClust^6^), to give a final set of 580 tumour-depth-CNA-SNV-subclonal reconstruction algorithm combinations. Each combination was evaluated using the scoring framework outlined above (**Figure 5, Supplementary Figure 4, Supplementary Tables 1,2**). In general, MuTect and SomaticSniper are more sensitive to low frequency variants and potentially preferable for subclonal reconstruction^46,47^. MuTect achieved the highest SNV-detection sensitivity in our synthetic tumours (mean sensitivity 0.65 ± 0.037 standard error), followed by Strelka (0.59 ± 0.032), SomaticSniper (0.50 ± 0.031) and finally MutationSeq (0.46 ± 0.045).

**Figure 5.**
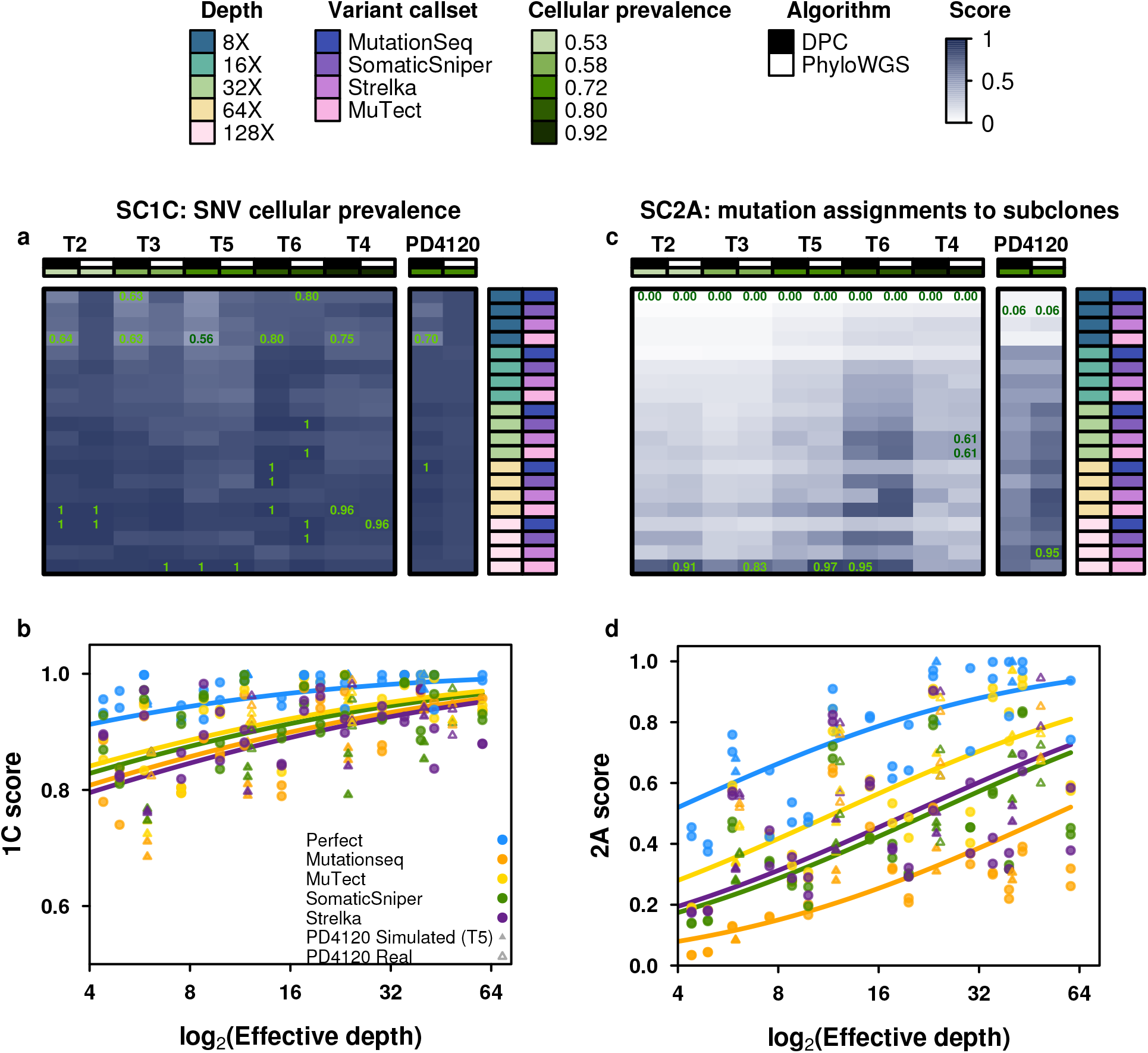
Error profiles of subclonal reconstruction algorithms. To identify general features of subclonal reconstruction algorithms, we created a set of tumour-depth-CNA-SNV-subclonal reconstruction algorithm combinations by using the framework outlined in Figure 3 and 4 to simulate five tumours with known subclonal architecture, followed by evaluation of two CNA detection approaches, five SNV detection methods, five read-depths and two subclonal reconstruction methods. The resulting reconstructions were scored using the scoring harness described in Figure 2, creating a dataset to explore general features of subclonal reconstruction methods. All scores are normalised to the score of the best performing algorithm when using perfect calls at the full tumour depth. Scores exceeding this baseline likely represent noise or overfitting and were capped at 1. **(a)** For SC1C (identification of the number of subclones and their cellular prevalence), all combinations of methods perform well. **(b)** By contrast, for SC2a (detection of the mutational characteristics of individual subclones), there is large inter-tumour variability in performance. **(c)** Score for SC1c (same as a) as a function of effective read-depth (depth after adjusting for purity and ploidy) improves with increased read-depth, and also changes with the somatic SNV detection method, with MuTect performing best, but still lagging perfect SNV calls by a significant margin. **(d)** Scores in SC2A show significant changes in performance as a function of effective read-depth.

This large-scale benchmarking of 580 simulated tumours reveals general features of subclonal reconstruction accuracy. For example, consider SC1C: estimation of SNV cellular prevalence. All algorithms and SNV detection algorithms showed a consistent increase in accuracy with increasing sequencing depth for SC1C (**Figure 5a, b**). No somatic SNV detection algorithm matched the performance of perfect SNV calls (β = 0.22, P = 0.0011, generalised linear model). By contrast, the use of high-*vs.* low-depth sequencing for subclonal detection of CNAs had no detectable influence on reconstruction accuracy in either real or simulated tumours (P>0.05; **Supplementary Table 2**). Interestingly, in SC1C, neither the use of low-*vs.* high-depth tumours for CNA detection nor the specific subclonal reconstruction algorithm used had a significant influence on the accuracy of subclonal reconstruction. Similarly, both PhyloWGS and DPClust performed interchangeably on this question in the simulated tumours (P=0.14, t=-1.47, **Supplementary Figure 5g-1, Supplementary Tables 2**).

A different story emerged for SC2A – identifying the mutational profiles of individual subclones (**Figure 5c,d**). All algorithms performed relatively poorly, with major intertumour differences in performance. Tumour T2 was systematically the most challenging to reconstruct and T6 the easiest (**Figure 5c, Supplementary Table 5**). This in part reflects the higher purity of T6, and indeed we see a strong association between effective read-depth and reconstruction accuracy in both the simulated and real tumours, with each additional doubling in read-depth increasing reconstruction score by about 0.1 (**Figure 5d**). At effective read-depths above 60x, the performance of all tumour-CNA-SNV-subclonal reconstruction combinations seemed to plateau, suggesting that a broad range of approaches can be effective for detection of subclonal mutational profiles at sufficient read-depth. Again, the use of high-vs. low-depth sequencing for subclonal CNA detection had no discernible influence (and this held true for all sub-challenges; **Supplementary Table 2**). By contrast, SC2A scores were strongly dependent on the SNV detection pipeline, with perfect calls out-performing the best individual algorithm (MuTect) by ~0.05 at any given read-depth. Differences in SNV detection algorithm sensitivity largely accounted for performance differences among algorithms (β_sensitivity_ = 0.30, P = 8.92 x 10^-13^, generalised linear models, **Supplementary Table 3**). MuTect, the most sensitive SNV detection algorithm, had the best performance and MutationSeq, the least sensitive, had the poorest. Broadly, SomaticSniper and Strelka showed similar performance, but interestingly showed significant tumour-by-algorithm interactions for several sub-challenges (**Supplementary Figure 5a-f**), which may reflect tumour-specific variability in their error profiles. Notably, MutationSeq performed much better on with the real tumour than with simulated tumours (**Supplementary Figure 5a-f**).

In general, DPClust and PhyloWGS showed very similar performance, but with exceptions that reflect their underlying algorithmic features. First, in SC1A DPClust, which uses purity measures derived from CNA reconstructions, showed a significant and systematic advantage over PhyloWGS (β_phyioWGS_ = −0.42, P = 1.5 x 10^-7^, generalised linear model), which uses purity measures partially dependent on SNV clustering. The latter are more sensitive to errors in VAF due to low sequencing depth and this is reflected in the pattern of SC1A scores. Second, in SC2A PhyloWGS, which uses a phylogenetically-aware clustering model, had significantly better performance than DPClust, which uses a flat clustering model (**Supplementary Figure 5g**). Thus, our metrics are sensitive to differences in modelling approaches, which manifest in variability in performance on different aspects of subclonal reconstruction. Validating these results, for the real high-depth tumour, DPClust significantly outperformed PhyloWGS in SC1, while PhyloWGS was superior in SC2 (**Supplementary Table 4**).

### Robustness of subclonal reconstruction to CNA detection errors

Surprised by the insensitivity of scores to the use of high- or low-depth sequencing data for subclonal CNA assessment, we sought to characterize the sensitivity of subclonal reconstruction to errors in CNA detection. We repeated the analyses described above using five types of CNA input: original (untouched), CNAs with doubled ploidy, CNA calls with a random portion of existing calls wrongly assigned (scramble) and CNAs with additional gains (scramble gains), or with additional losses (scramble loss). The latter three error types were titrated in intensity, scrambling 10%, 20%, 30%, 40% and 50% of all CNAs, gains and losses, respectively.

The resulting 4,250 tumour-depth-CNA-SNV-reconstruction combinations were each assessed using our scoring metrics (**Supplementary Table 1**). For SC1 and SC2, incorrect ploidy impaired reconstruction accuracy overall (**Figure 6A**). As expected, scores decreased as the proportion of incorrectly assigned CNAs increased (**Supplementary Figure 6a,b**). The effect of incorrect calls on SC2A accuracy was only apparent at >32x coverage and was strongest with perfect and MuTect SNVs (**Figure 6B**), suggesting the relative impact of CNA errors increases with reconstruction quality. Interestingly, PhyloWGS had significantly better performance for all subchallenges than DPClust when CNA errors were introduced (SC1C: βphyiowos = 0.042, P = 6.06 x 10^-10^; SC2A: β _PhyloW_GS = 0.066, P = 1.85 x 10^-10^ generalised linear models; **Supplementary Table 5**). These results suggest that PhyloWGS’s strategy of incorporating CNAs in the allele count model may be more robust to errors in CNA detection than only using them to initially correct SNV VAFs (**Supplementary Figure 5g, Online methods**). As CNA-handling in the presence of errors distinguishes algorithms with otherwise comparable performance, increasing robustness to errors in CNA calls may be a promising avenue for improvement of subclonal reconstruction algorithms.

**Figure 6.**
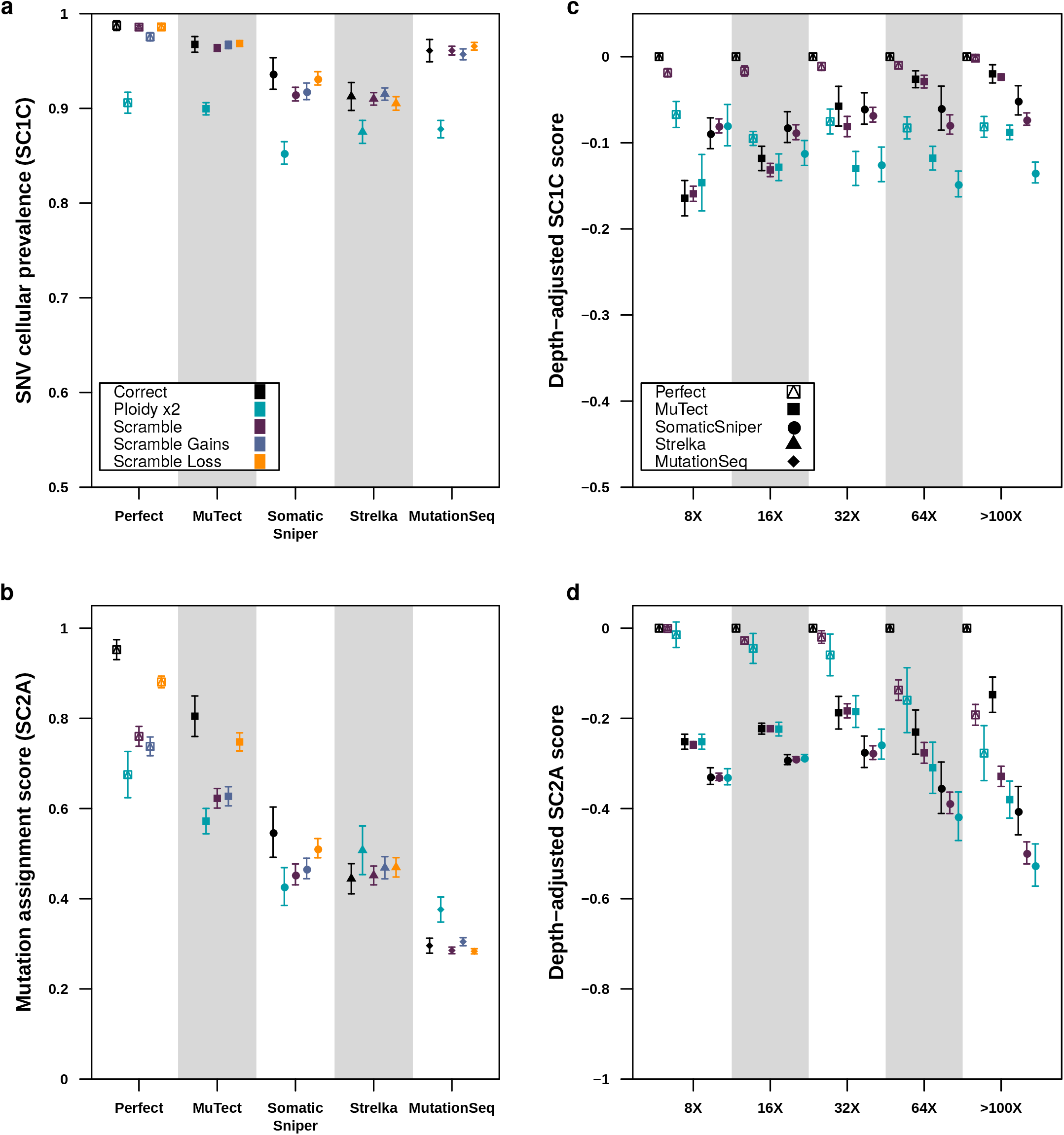
Impact of CNA error profiles on subclonal reconstruction. **(a)** Effect of CNA errors on mean SC1c scores and SC2a **(b)** scores (with standard errors shown) at 100x across somatic SNV detection algorithms. **(c)** Effect of CNA errors on mean SC1c and SC2a **(d)** scores (with standard errors shown) at various depths when scores for perfect calls are set to zero to yield depth-adjusted scores.

Taken together, these results suggest that subclonal reconstruction accuracy is highly sensitive both to SNV and CNA detection, with interactions between specific pairs of variant detection and subclonal reconstruction algorithms(**Online methods; Supplementary Figure 6c,d**). There is significant room for algorithmic improvements that capture inter-tumour differences and better model the error characteristics of feature-detection pipelines.

## Discussion

Increasingly large numbers of tumours receive genomic interrogation each year as DNA sequencing costs diminish and evidence for clinical utility increases. Nevertheless, it remains common practice for only a single spatial region of a cancer to be sequenced. The reasons for this are myriad: costs of multi-region sequencing, needs to preserve tumour tissue for future clinical use and increasing analysis of scarce biopsy-derived specimens in diagnostic and metastatic settings. Whilst robust subclonal reconstruction from multi-region sequencing is well-known^5–8^, accurately reconstructing tumour evolutionary properties from single-region sequencing could open new avenues for linking these to clinical phenotypes and outcomes.

We describe a framework for evaluating single-sample subclonal reconstruction methods, comprising a novel way of scoring their accuracy, a technique for phasing short-read sequencing data, an enhanced read-level simulator of tumour genomes with realistic biological properties and a portable software framework for rapidly and consistently executing a library of subclonal reconstruction algorithms. These elements, each implemented as open-source software and independently reusable, form an integrated system for quantitation of key parameters of subclonal reconstruction. We generate a 580-tumour titration-series for evaluating subclonal reconstruction sensitivity to both effective read depth and specific somatic SNV detection pipelines. These data give guidance for improving subclonal reconstruction: increasing effective read-depth above 60x, after controlling for tumour purity and ploidy. They also suggest reconstruction algorithm developers should consider accounting for the error properties of specific somatic variant detection approaches.

Lineage-tracing tools are emerging that will likely revolutionize our understanding of tissue growth and evolution, such as GESTALT^48^, ScarTrace^49^, and MEMOIR^50^. However, these are not applicable to the study of human cancer tissues *in vivo*. In many areas of biology, ground-truth is still either inaccessible or impractical to measure with precision. In cases like these, simulations are extremely valuable in providing a lower bound on error profiles and an upper bound on method accuracy. By incorporating all currently known features of a phenomenon, simulators codify our understanding. Divergence between simulated and real results quantitates the gaps in our knowledge. The creation of an open-source, freely available simulator capturing most known features of cancer genomes thus represents one avenue for exploring the boundaries of our knowledge.

Large-scale benchmarking of multiple subclonal reconstruction methods using this framework on larger numbers of tumours is needed to create a gold-standard. Such a benchmark would both inform algorithm users, who will benefit from an understanding of the specific error profiles of different methods, and algorithm developers who will be able to update and improve methods while ensuring software portability. Tumour simulation frameworks provide a valuable way for method benchmarking, and can complement other approaches like comparison of single-region to multi-region subclonal reconstruction, and the use of model organism and sample-mixing experiments.

## Accession Codes

Sequences files are available at EGA under study accession number EGAS00001002092. BAMSurgeon is available at: https://github.com/adamewing/bamsurgeon. The framework for subclonal mutation simulation is available at: http://search.cpan.org/~boutroslb/NGS-Tools-BAMSurgeonv1.0.0/. The PhaseTools BAM phasing toolkit is available at https://github.com/mateidavid/phase-tools. Scripts providing the complete scoring harness are available at: https://github.com/Sage-Bionetworks/SMC-Het-Challenge.

## Acknowledgements

The authors thank the members of their labs for support, and Sage Bionetworks and the DREAM Challenge organization for their ongoing support of the SMC-Het Challenge. In particular, they thank T. Norman, J.C. Bare, S. Friend and G. Stolovitzky for their patience, technical support and scientific insight. The authors also thank Ruping Sun and Christina Curtis for kindly sharing code for calculating the intra-tumour heterogeneity metrics and building the support vector machine predictor in multi-region sequencing simulations. This study was conducted with the support of the Ontario Institute for Cancer Research to P.C.B. and J.T.S. through funding provided by the Government of Ontario. This work was supported by Prostate Cancer Canada and is proudly funded by the Movember Foundation – Grant #RS2014-01. This study was conducted with the support of Movember funds through Prostate Cancer Canada and with the additional support of the Ontario Institute for Cancer Research, funded by the Government of Ontario. This project was supported by Genome Canada through a Large-Scale Applied Project contract to P.C.B., S.P. Shah and R.D. Morin. This work was supported by the Discovery Frontiers: Advancing Big Data Science in Genomics Research program, which is jointly funded by the Natural Sciences and Engineering Research Council (NSERC) of Canada, the Canadian Institutes of Health Research (CIHR), Genome Canada and the Canada Foundation for Innovation (CFI). This work was supported by an NSERC operating grant grant and a gift from the NVIDIA Foundation, an advised fund of Silicon Valley Community Foundation, to Q.D.M. This research is part of the University of Toronto’s Medicine by Design initiative, which receives funding from the Canada First Research Excellence Fund (CFREF). J.A.W. was partially supported by an Ontario Graduate Scholarship. This work was supported by the Francis Crick Institute, which receives its core funding from Cancer Research UK (FC001202), the UK Medical Research Council (FC001202), and the Wellcome Trust (FC001202). M.T. is a postdoctoral fellow supported by the European Union’s Horizon 2020 research and innovation program (Marie Skłodowska-Curie Grant Agreement No. 747852-SIOMICS). P.V.L. is a Winton Group Leader in recognition of the Winton Charitable Foundation’s support towards the establishment of The Francis Crick Institute. P.C.B. was supported by a Terry Fox Research Institute New Investigator Award and a CIHR New Investigator Award. D.C.W. is supported by the Li Ka Shing foundation. The Galaxy portions of the evaluation system were supported by NIH Grants U41 HG006620 and R01 AI134384-01 as well as NSF Grant 1661497. The following NIH grants supported this work: R01-CA180778 (J.M.S.) and U24-CA143858 (J.M.S.). The authors thank Google Inc. (in particular N. Deflaux) for their ongoing support of the ICGC-TCGA DREAM Somatic Mutation Calling Challenge. This work was supported by the NIH/NCI under award number P30CA016042.

## Author contributions

All authors: Edited & approved final manuscript. A.S. wrote first draft of paper, designed experiments performed statistical analyses, performed bioinformatics analyses, performed data visualisation. M.T. wrote first draft of paper, designed experiments, generated tools & reagents, performed statistical analyses, performed bioinformatics analyses, performed data visualisation. S.M.G.E. wrote first draft of paper, generated tools & reagents, performed bioinformatics analyses, performed data visualisation. A.G.D. wrote first draft of paper, designed experiments, generated tools & reagents, performed bioinformatics analyses. M.D. generated tools & reagents. S.D. generated tools & reagents. L.Y.L. generated tools & reagents. S. S. generated tools & reagents. H.Z. generated tools & reagents. K.Z. generated tools & reagents, performed bioinformatics analyses. T.O.Y. generated tools & reagents, performed bioinformatics analyses. J.M.C. generated tools & reagents. A.B. generated tools & reagents. C.M.L. generated tools & reagents. I.U. generated tools & reagents. B.L. generated tools & reagents. A.D.E. generated tools & reagents, supervised research. NMW performed bioinformatics analyses, performed data visualisation. J.A.W. performed bioinformatics analyses. M.K.H.Z. performed bioinformatics analyses. C.V.A. performed bioinformatics analyses. C.P. performed data visualisation. J.T.S. supervised research. J.M.S. supervised research. D.A. supervised research. Y.G. supervised research. K.E. wrote first draft of paper, supervised research. D.C.W. designed experiments, supervised research. Q.D.M. wrote first draft of paper, designed experiments, generated tools & reagents, supervised research. P.V.L. wrote first draft of paper, designed experiments, supervised research. P.C.B. wrote first draft of paper, designed experiments, supervised research.

## Supplementary Figure Legends

**Supplementary Figure 1 | Behaviour of Scoring Metrics**

**(a)** The score for each candidate mutation cluster assignment score (SC2) metric considered with an increasing proportion of mutations assigned to the wrong useful clusters. **(b)** The score for each candidate SC2 metric considered with an increasing proportion of mutations in noise clusters. **(c)** The score for each candidate SC2 metric considered as the number of predicted clusters increases. The true number of clusters (four) is marked by the vertical line. Excess clusters retain correct co-clustering and are subsets of the true clusters. **(d)** For each potential SC2 scoring metric, the proportion of simulation runs that satisfied each of the four desirable metric properties for a given simulation parameter setting. Each property is tested by fixing all but one of the simulation parameters and then looking at the effect of changing the fourth parameter on the metric score.

**Supplementary Figure 2 | Mutation Phasing**

**(a)** Example of the PhaseTools algorithm constructing an extended phase set from four heterozygous sites by leveraging NGS and parent phasing. ngs_phasing of 5 heterozygous sites in the child and the corresponding NGS-phased sites in the mother and father, shown with informative NGS fragments. Heterozygote variants boxed together represent phase sets. There is not enough information to construct a single phase set. **(b)** parent_base phasing uses parental genotypes to assign parent of origin to the 5 hets in the child. Hets 2 and 5 remain unresolved while heterozygotes 1, 3, and 4 show at least one unambiguous parent of origin. **(c)** parent_ngs phasing extends parent_base phasing with parental NGS fragments from ngs_phasing. The linked NGS fragment in sites 2 and 3 (T, T) of the maternal genotype is not informative as site 3 is homozygous, however the linked NGS fragment in sites 2 and 3 (T and A) of the paternal genotype is heterozygous and therefore informative. The phasing proposed by ngs_father of sites 3 and 4 (GG/AC) contradicts parent of origin information in hets 3 and 4 (A and G). This event is recognised as a pre-meiosis recombination event in the child and the ngs_father phasing is ignored. **(d)** ngs+parent_ngs phasing extends ngs_child phasing with parent_ ngs, giving priority to ngs_child phasing. NGS fragments such as hets 2 and 3 (T and T) take precedent over any phasing assigned by parent_ngs phasing see hets 2 and 3 (C and T) and indicate probable recombination events (shown with diagonal lines). Two possible sets of recombination events are shown. The proximity between phased heterozygotes determines which recombination events are most probable. Here, the recombination events shown on the right are selected, as recombination between sites 1 and 2 is more likely than recombination between sites 3 and 4, as sites 1 and 2 are further apart. The final phase sets are shown. **(e)** Schematic of phase-set reconstruction. Priority is given to procedures on the left. **(f)** Genome-wide logR and BAF tracks from a simulated BAM using the original “naive” BAMSurgeon. **(g)** Same as (a) with the new proposed BAMSurgeon pipeline and additional whole-chromosomal events. **(h)** Comparison of expected and observed VAFs for SNVs on chromosome 17 of a tumour simulated with the “naive” BAMSurgeon (top panel) and the new pipeline (bottom panel) with Pearson correlations. As naive BAMSurgeon does not simulate allele-specific gains and losses, copy number alterations in one allele do not produce the expected allele-frequency changes for each phase. Tumour structure showing a clonal deletion and duplication on the chromosome is specified in the top right panel. Allele-specific copy number events for phase B (phase A has no copy number alterations) and the subclone where they occur are shown in the top heatmap. Expected VAFs were calculated by summing CNA-adjusted CCF of the leaf subclones were the SNV was present, adjusting for the frequency of the SNV in those subclones and standardizing by the total CNA-adjusted depth *n* that region. **(i)** Independent component analysis of the intra-tumour heterogeneity metrics presented in Sun et al. on 1,366 3D simulated training tumours (Sun et al. simulator) and 5×3-region BAMs derived from 5 simulated 3D test tumours (Sun *et al.* simulator) using BAMSurgeon. The 5 regions were classified correctly using an SVM predictor (**Online Methods**).

**Supplementary Figure 3 | Real and Simulated Subclonal Structures**

True subclonal structures of the simulated tumours (**T2, T3, T4, T5** and **T6**) that were simulated with their desired and observed variant allele frequency histograms and logR profiles. In each panel, we show the phylogenetic tree, inspired by published reconstructed tumours, and the mutations associated with each (sub)clone. The top figures compared expected cancer cell fractions of the SNVs under a diploid setting, against the inferred cancer cell fractions from the simulated data. T5, for which the inferred purity is over-estimated due to the limitations of the copy number detection algorithm to identify subclonal whole genome duplication, shows an observed space that departs from the expected. The bottom figures compare the observed and expected BAF and logR of the genomic segments identified by the copy number detection algorithm.

**Supplementary Figure 4 | Subclonal Reconstruction Scores**

Subclonal reconstruction scores based on the five tumours with each somatic SNV detection algorithm-depth-algorithm combination. All scores are normalised to the score of the best performing algorithm using perfect calls at the full tumour depth. Scores exceeding this baseline likely represent noise or overfitting and were capped at 1. **(a)** Scores for 1A are uniformly high. **(b)** Scores for SC1B improve with depth and but not continuous as the metric reflects a true proportion. **(c)** Clonal fraction is low below 32X but rapidly increases at higher depths, with some inter-tumour variability. **(d,f)** Scores for SC2B **(d)** and SC3B **(f)** closely mirror those of SC2A and SC3A **(e)**, respectively.

**Supplementary Figure 5 | Covariates of Scoring Accuracy**

SC2A score increases with effective depth for all tumours but the effect of the somatic SNV detection algorithm depends on the tumour. (a) T2 (b) T3 (c) T4 (d) T5 (e) T6 (f) real PD4120. (g) A summary of the differences between DPClust and PhyloWGS. PhyloWGS incorporates CNAs and phylogenetic structure into the generative model sampled through Markov chain Monte-Carlo (MCMC). (h-l) Comparison of subclonal reconstruction scores for each sub-challenge using PhyloWGS (x-axis) and DPClust (y-axis). Somatic SNV detection algorithms are coded by colour and tumours are coded by symbol. P-values and effect size for a t-test between algorithms on score are shown.

**Supplementary Figure 6 | Effects of Noise in CNA Detection on Reconstruction Accuracy**

Effect of proportion of CNAs scrambled on a representative tumour (T2) at 128x sequencing coverage. **(a)** Allele specific copy number profiles as increasing proportions of CNAs are randomly assigned to a wrong copy number state. **(b)** Effect of increasing the proportion of wrongly assigned CNAs on SC2a scores at 128x for different somatic SNV detection algorithms and SRC algorithms. **(c)** Deriving an imperfect “truth” from real data and scoring runs against it. We took data from 538 donors from the PCAWG study^41^ and executed DPClust on different sets of SNVs: a consensus set of SNVs was used as “truth” and three individual somatic SNV detection pipelines (MuTect, DKFZ and Sanger on mutations in a (C>T)pG context). We also executed PhyloWGS on the consensus sets. We then scored SC1C for each run against the “truth”. **(d)** We took data from 10 donors from the PCAWG study with at least 5 tumour regions or metastases sequenced and ran DPClust in 5 dimensions to derive a “truth” from it. We scored each one-dimensional DPClust run on individual region against the “truth” derived from the multi-dimensional run for SC1C and SC2A and compared them against scores obtained from randomised 1C and 2A inputs (**Online Methods**).

