## Supplementary Figures for "Creating Standards for Evaluating Tumour Subclonal Reconstruction"

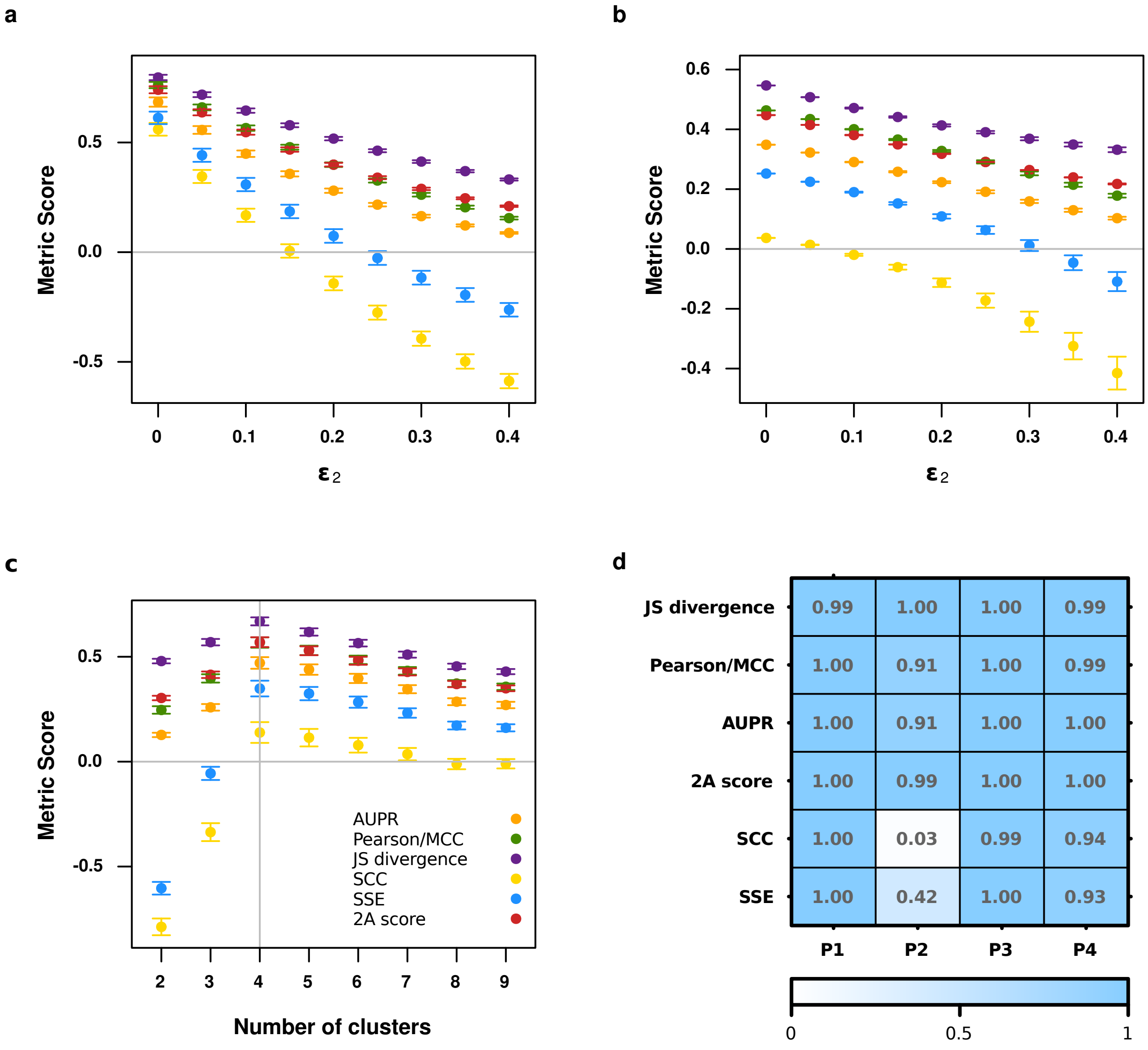


Supplementary Figure 1

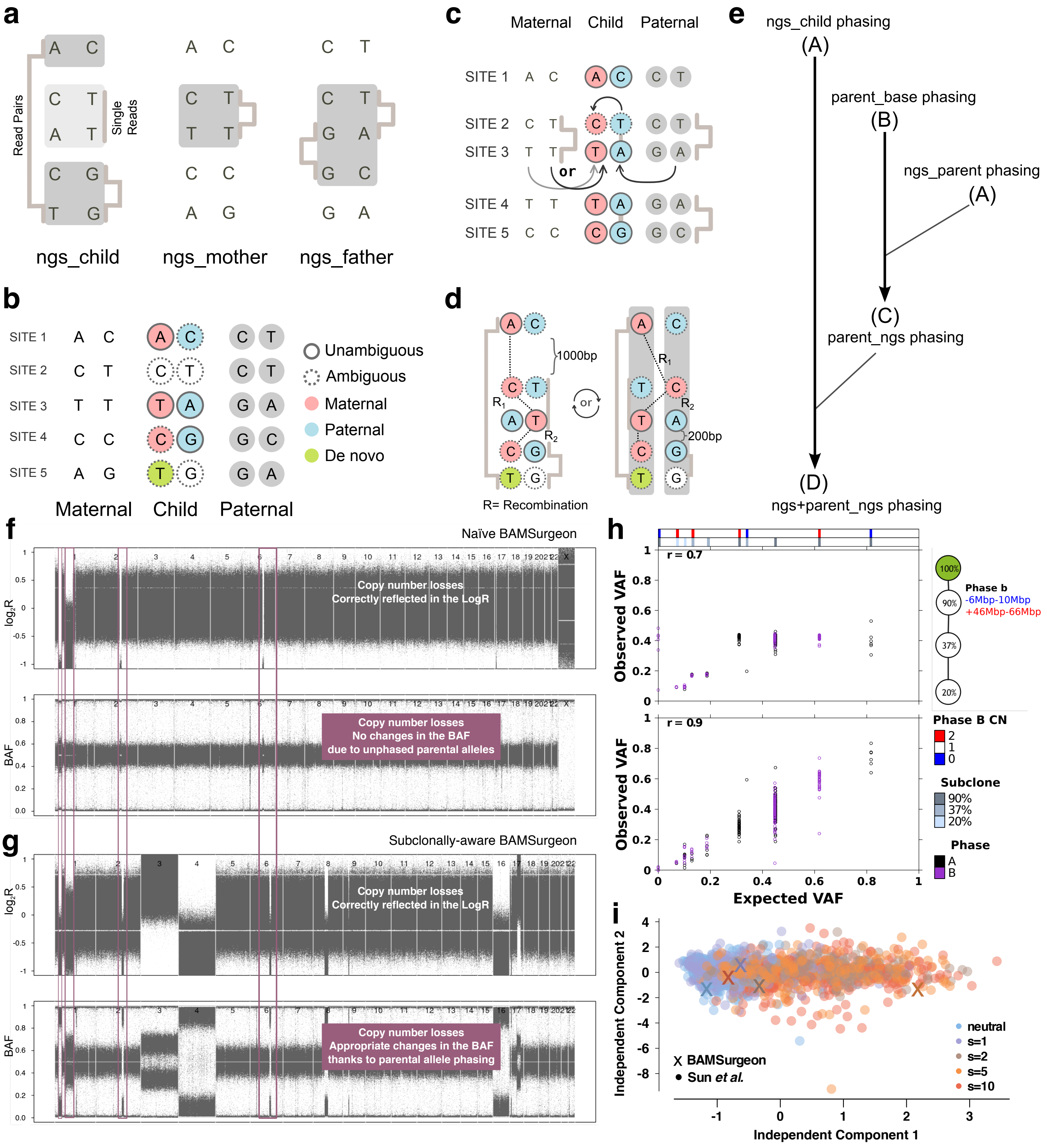


Supplementary Figure 2

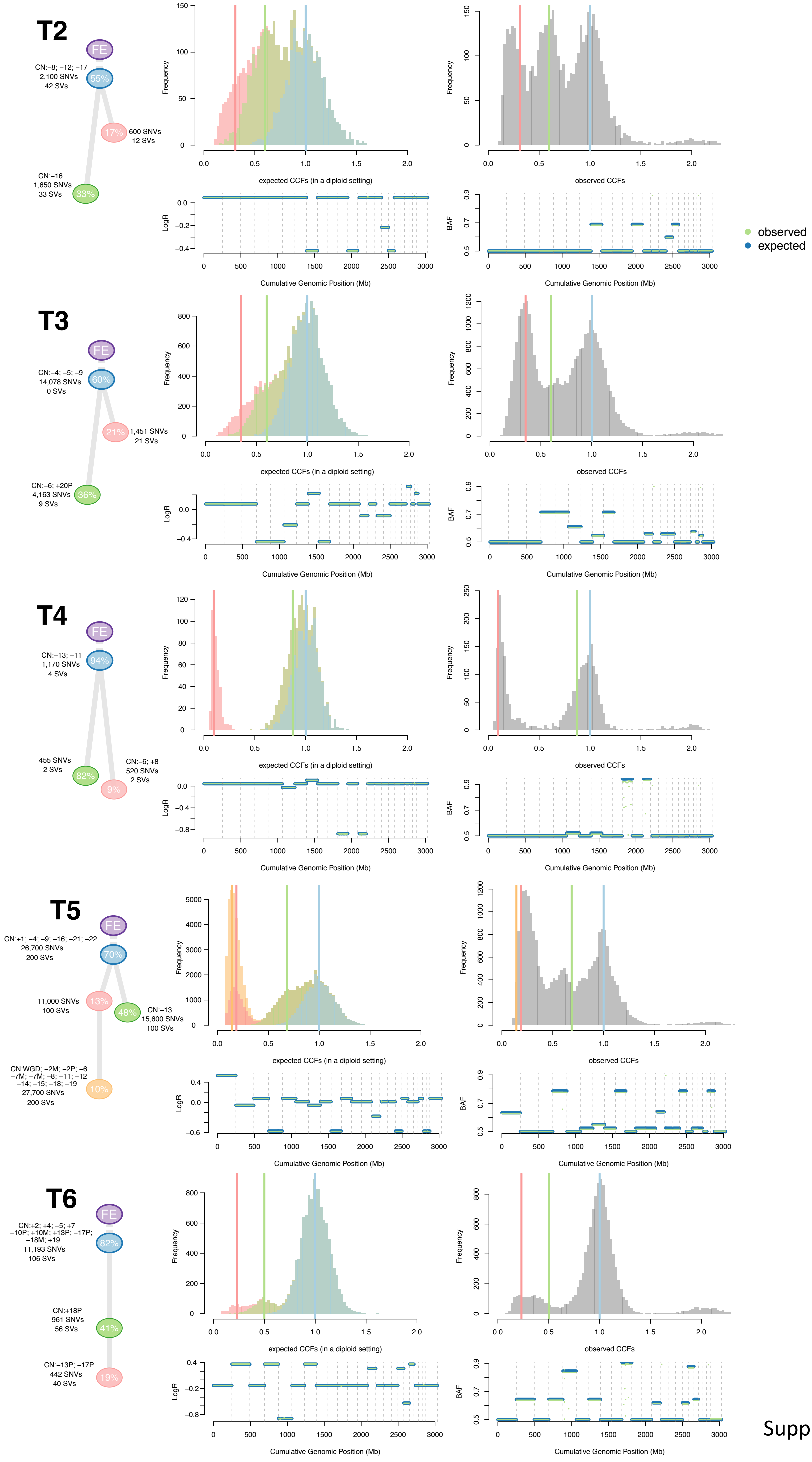


Supplementary Figure 3

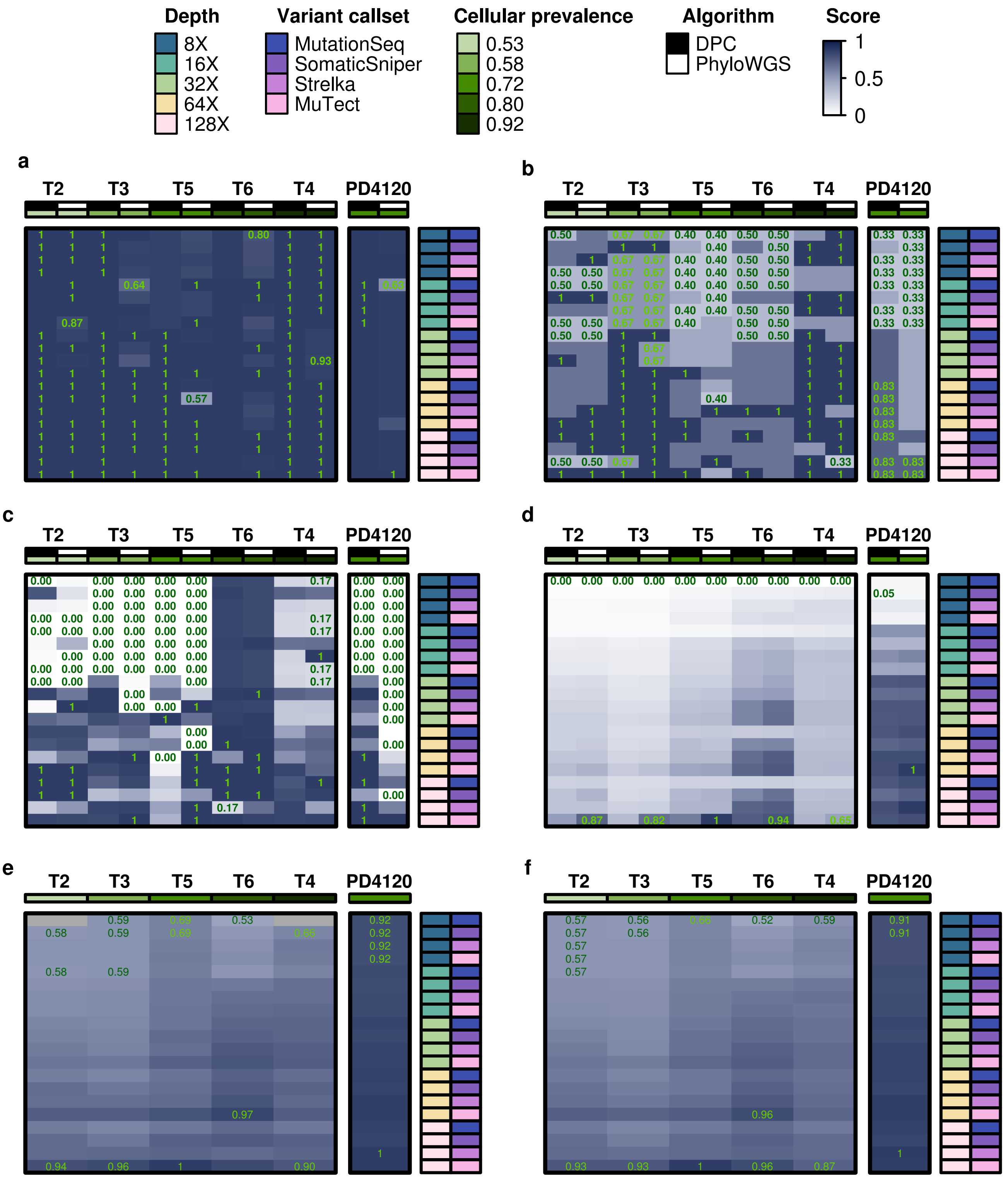


Supplementary Figure 4

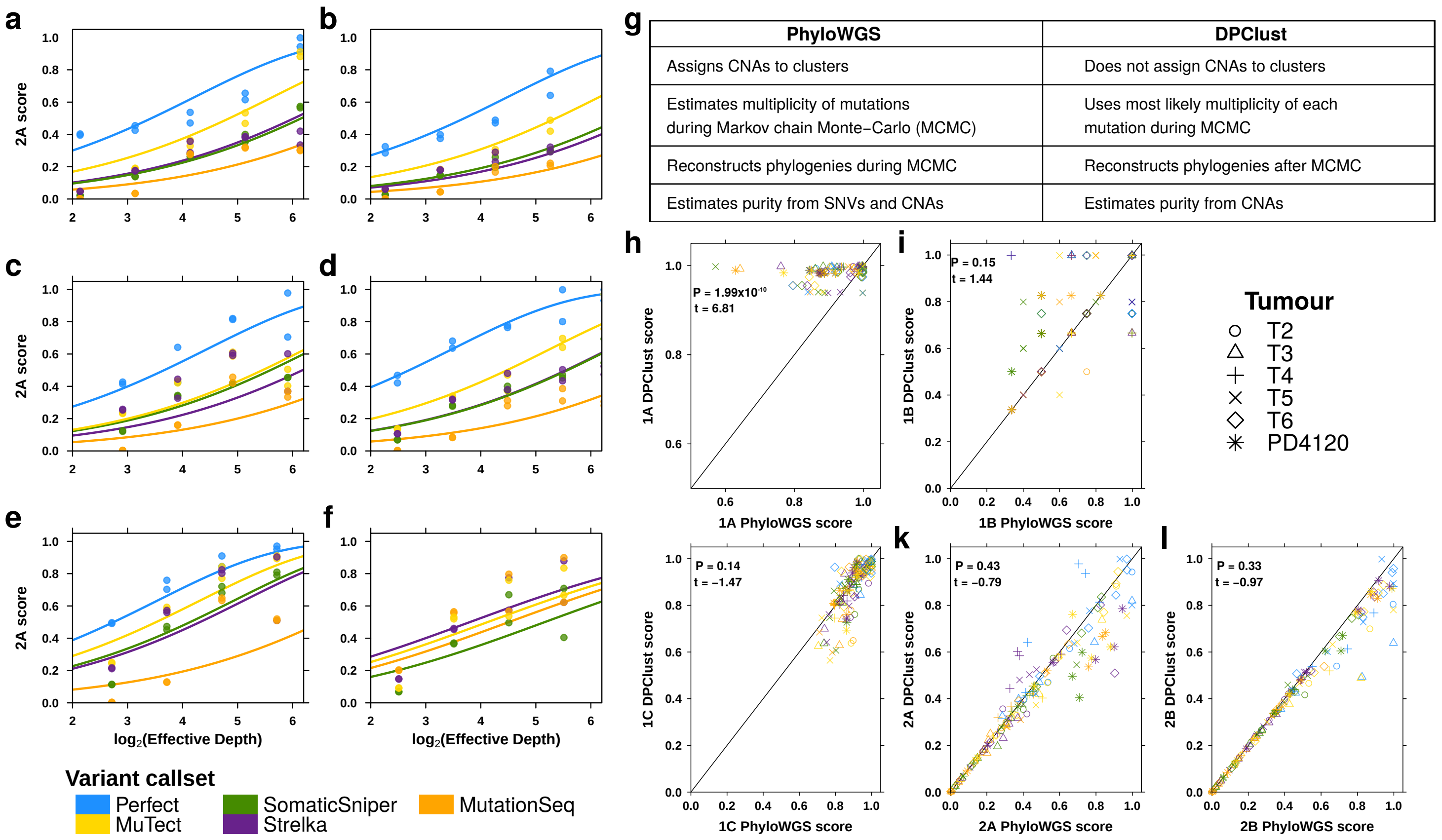


| PhyloWGS | DPClust |
|---|---|
| Assigns CNAs to clusters | Does not assign CNAs to clusters |
| Estimates multiplicity of mutations during Markov chain Monte−Carlo (MCMC) | Uses most likely multiplicity of each mutation during MCMC |
| Reconstructs phylogenies during MCMC | Reconstructs phylogenies after MCMC |
| Estimates purity from SNVs and CNAs | Estimates purity from CNAs |

Supplementary Figure 5

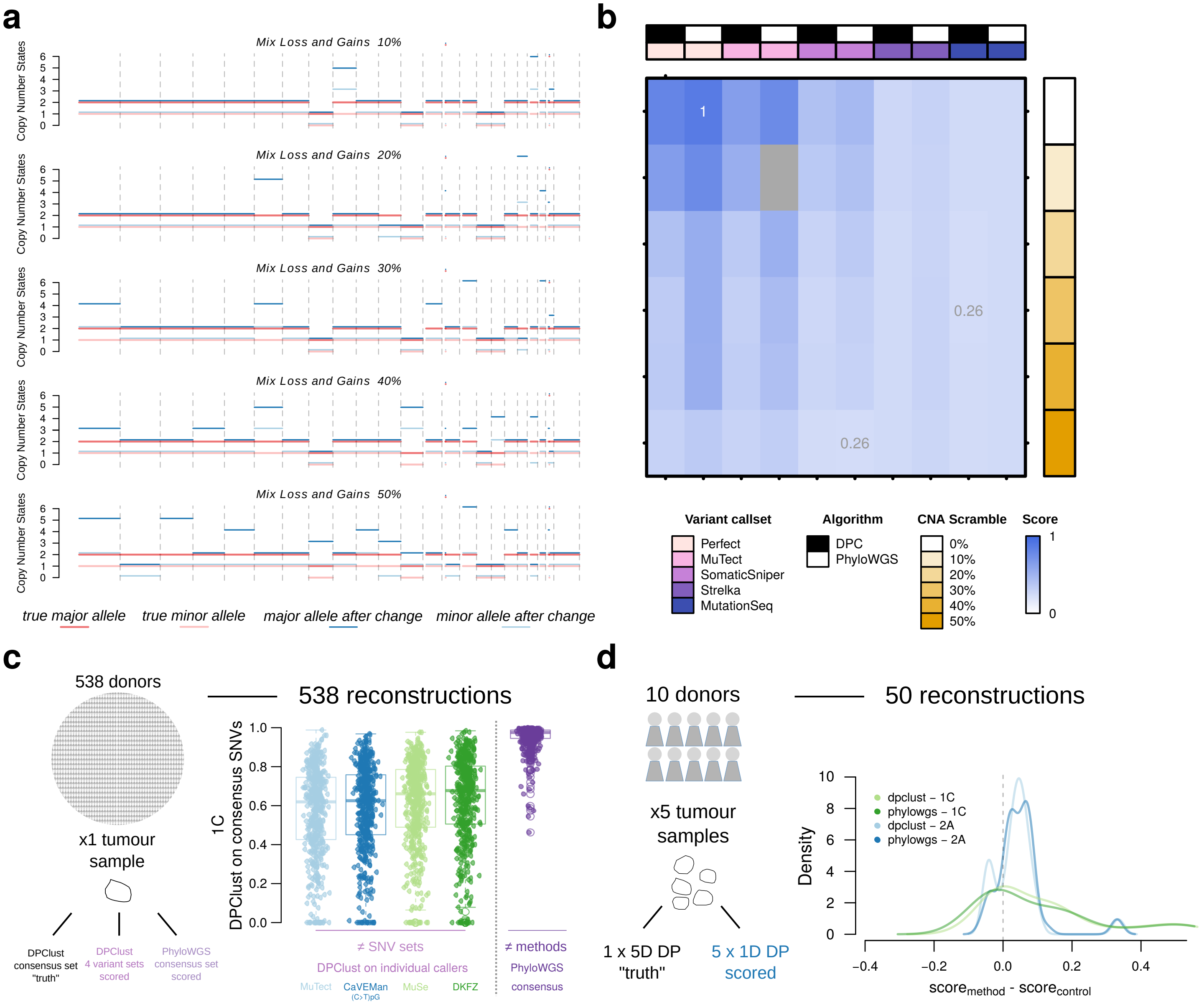


Supplementary Figure 6
