## Supplementary Table 2 for "Creating Standards for Evaluating Tumour Subclonal Reconstruction"

Supplementary Table 2: Genaralized linear models for each subchallenge. $\beta$ regressions were used for all subchallenges except 1B where a binomial regression was used. N=500 for subchallenges 1 and 2 and N=250 for subchallenge 3.

| $\beta$ | Estimate 1A | Std. Error 1A | P-value 1A | Estimate 1B | Std. Error 1B | P-value 1B | Estimate 1C | Std. Error 1C | P-value 1C | Estimate 2A | Std. Error 2A | P-value 2A | Estimate 2B | Std. Error 2B | P-value 2B | Estimate 3A | Std. Error 3A | P-value 3A | Estimate 3B | Std. Error 3B | P-value 3B |
|---|---|---|---|---|---|---|---|---|---|---|---|---|---|---|---|---|---|---|---|---|---|
| Intercept | 4.42 | 0.15 | $1.30 \times 10^{-196}$ | 1.76 | 0.13 | $3.73 \times 10^{-39}$ | 1.23 | 0.05 | $3.77 \times 10^{-133}$ | 0.46 | 0.08 | $6.91 \times 10^{-9}$ | 0.34 | 0.07 | $5.54 \times 10^{-6}$ | 1.30 | 0.08 | $1.18 \times 10^{-61}$ | 1.37 | 0.08 | $6.39 \times 10^{-71}$ |
| Effective Depth | 0.25 | 0.04 | $3.78 \times 10^{-10}$ | 0.34 | 0.09 | $1.14 \times 10^{-4}$ | 0.25 | 0.01 | $6.08 \times 10^{-147}$ | 0.56 | 0.02 | $4.72 \times 10^{-254}$ | 0.58 | 0.02 | $1.78 \times 10^{-294}$ | 0.31 | 0.01 | $1.52 \times 10^{-98}$ | 0.31 | 0.01 | $2.43 \times 10^{-118}$ |
| T3 | -0.16 | 0.13 | $2.14 \times 10^{-1}$ | -0.26 | 0.13 | $3.97 \times 10^{-2}$ | -0.02 | 0.07 | $8.23 \times 10^{-1}$ | -0.11 | 0.11 | $2.90 \times 10^{-1}$ | -0.14 | 0.10 | $1.60 \times 10^{-1}$ | -0.00 | 0.11 | $9.94 \times 10^{-1}$ | -0.06 | 0.11 | $5.50 \times 10^{-1}$ |
| T4 | -0.04 | 0.13 | $7.66 \times 10^{-1}$ | -0.42 | 0.13 | $9.45 \times 10^{-4}$ | -0.22 | 0.07 | $1.45 \times 10^{-3}$ | -0.00 | 0.11 | $9.81 \times 10^{-1}$ | 0.02 | 0.10 | $8.45 \times 10^{-1}$ | 0.09 | 0.11 | $4.27 \times 10^{-1}$ | -0.02 | 0.11 | $8.76 \times 10^{-1}$ |
| T5 | -0.44 | 0.13 | $5.15 \times 10^{-4}$ | -0.62 | 0.12 | $1.34 \times 10^{-7}$ | -0.01 | 0.07 | $9.09 \times 10^{-1}$ | 0.37 | 0.11 | $6.53 \times 10^{-4}$ | 0.45 | 0.10 | $1.80 \times 10^{-5}$ | 0.41 | 0.12 | $6.35 \times 10^{-4}$ | 0.34 | 0.12 | $3.65 \times 10^{-3}$ |
| T6 | -0.72 | 0.12 | $5.26 \times 10^{-9}$ | -0.37 | 0.13 | $3.69 \times 10^{-3}$ | 0.07 | 0.07 | $2.98 \times 10^{-1}$ | 0.48 | 0.11 | $1.36 \times 10^{-5}$ | 0.63 | 0.11 | $1.89 \times 10^{-9}$ | 0.50 | 0.12 | $4.55 \times 10^{-5}$ | 0.41 | 0.12 | $5.79 \times 10^{-4}$ |
| PhyloWGS | -0.42 | 0.08 | $1.47 \times 10^{-7}$ | -0.09 | 0.08 | $2.64 \times 10^{-1}$ | 0.07 | 0.02 | $5.38 \times 10^{-4}$ | 0.05 | 0.03 | $1.28 \times 10^{-1}$ | 0.14 | 0.03 | $1.25 \times 10^{-6}$ | | | | | | |
| Downsampling | -0.18 | 0.08 | $2.20 \times 10^{-2}$ | 0.07 | 0.08 | $3.67 \times 10^{-1}$ | -0.02 | 0.02 | $3.03 \times 10^{-1}$ | 0.00 | 0.03 | $9.97 \times 10^{-1}$ | -0.00 | 0.03 | $9.49 \times 10^{-1}$ | -0.01 | 0.03 | $8.25 \times 10^{-1}$ | -0.02 | 0.03 | $4.39 \times 10^{-1}$ |
| MuTect | -0.10 | 0.13 | $4.28 \times 10^{-1}$ | -0.33 | 0.12 | $8.46 \times 10^{-3}$ | -0.23 | 0.07 | $5.87 \times 10^{-4}$ | -0.63 | 0.11 | $3.38 \times 10^{-9}$ | -0.66 | 0.10 | $4.87 \times 10^{-11}$ | -0.41 | 0.10 | $6.77 \times 10^{-5}$ | -0.48 | 0.10 | $1.35 \times 10^{-6}$ |
| SomaticSniper | -0.11 | 0.13 | $4.03 \times 10^{-1}$ | -0.42 | 0.15 | $4.47 \times 10^{-3}$ | -0.22 | 0.07 | $1.14 \times 10^{-3}$ | -0.99 | 0.11 | $7.96 \times 10^{-20}$ | -1.02 | 0.10 | $3.34 \times 10^{-23}$ | -0.67 | 0.10 | $1.86 \times 10^{-11}$ | -0.74 | 0.10 | $1.78 \times 10^{-14}$ |
| Strelka | -0.25 | 0.12 | $4.16 \times 10^{-2}$ | -0.59 | 0.12 | $3.52 \times 10^{-7}$ | -0.32 | 0.07 | $2.90 \times 10^{-6}$ | -1.02 | 0.11 | $5.60 \times 10^{-21}$ | -0.96 | 0.10 | $1.34 \times 10^{-20}$ | -0.67 | 0.10 | $2.34 \times 10^{-11}$ | -0.69 | 0.10 | $9.35 \times 10^{-13}$ |
| Mutationseq | -0.13 | 0.13 | $2.86 \times 10^{-1}$ | -0.65 | 0.12 | $3.00 \times 10^{-8}$ | -0.30 | 0.07 | $1.08 \times 10^{-5}$ | -1.37 | 0.11 | $4.20 \times 10^{-35}$ | -1.30 | 0.11 | $3.38 \times 10^{-35}$ | -0.88 | 0.10 | $5.68 \times 10^{-17}$ | -0.85 | 0.10 | $7.52 \times 10^{-19}$ |
| MuTect:T3 | | | | | | | -0.13 | 0.10 | $1.82 \times 10^{-1}$ | -0.05 | 0.15 | $7.62 \times 10^{-1}$ | -0.08 | 0.14 | $5.62 \times 10^{-1}$ | -0.13 | 0.14 | $3.58 \times 10^{-1}$ | -0.12 | 0.14 | $4.03 \times 10^{-1}$ |
| MuTect:T4 | | | | | | | -0.18 | 0.10 | $6.09 \times 10^{-2}$ | -0.12 | 0.15 | $4.16 \times 10^{-1}$ | -0.09 | 0.14 | $5.20 \times 10^{-1}$ | -0.13 | 0.15 | $3.78 \times 10^{-1}$ | -0.13 | 0.14 | $3.56 \times 10^{-1}$ |
| MuTect:T5 | | | | | | | -0.21 | 0.10 | $2.63 \times 10^{-2}$ | -0.23 | 0.15 | $1.37 \times 10^{-1}$ | -0.12 | 0.14 | $4.25 \times 10^{-1}$ | -0.20 | 0.16 | $2.06 \times 10^{-1}$ | -0.15 | 0.15 | $3.16 \times 10^{-1}$ |
| MuTect:T6 | | | | | | | 0.02 | 0.10 | $8.21 \times 10^{-1}$ | 0.17 | 0.15 | $2.63 \times 10^{-1}$ | 0.09 | 0.15 | $5.42 \times 10^{-1}$ | -0.33 | 0.16 | $3.61 \times 10^{-2}$ | -0.24 | 0.15 | $1.18 \times 10^{-1}$ |
| SomaticSniper:T3 | | | | | | | -0.39 | 0.10 | $4.41 \times 10^{-5}$ | -0.18 | 0.15 | $2.45 \times 10^{-1}$ | -0.17 | 0.15 | $2.56 \times 10^{-1}$ | -0.15 | 0.14 | $2.88 \times 10^{-1}$ | -0.13 | 0.14 | $3.51 \times 10^{-1}$ |
| SomaticSniper:T4 | | | | | | | -0.04 | 0.10 | $6.61 \times 10^{-1}$ | -0.06 | 0.15 | $6.84 \times 10^{-1}$ | 0.03 | 0.15 | $8.62 \times 10^{-1}$ | -0.02 | 0.15 | $8.66 \times 10^{-1}$ | -0.03 | 0.14 | $8.56 \times 10^{-1}$ |
| SomaticSniper:T5 | | | | | | | -0.55 | 0.10 | $8.03 \times 10^{-9}$ | -0.26 | 0.15 | $8.70 \times 10^{-2}$ | -0.32 | 0.15 | $2.82 \times 10^{-2}$ | -0.30 | 0.15 | $5.04 \times 10^{-2}$ | -0.28 | 0.15 | $5.47 \times 10^{-2}$ |
| SomaticSniper:T6 | | | | | | | 0.02 | 0.10 | $8.57 \times 10^{-1}$ | 0.20 | 0.15 | $1.99 \times 10^{-1}$ | 0.09 | 0.15 | $5.49 \times 10^{-1}$ | -0.35 | 0.15 | $2.13 \times 10^{-2}$ | -0.27 | 0.15 | $6.33 \times 10^{-2}$ |
| Strelka:T3 | | | | | | | -0.10 | 0.10 | $2.85 \times 10^{-1}$ | -0.05 | 0.15 | $7.69 \times 10^{-1}$ | -0.07 | 0.15 | $6.10 \times 10^{-1}$ | -0.08 | 0.14 | $5.89 \times 10^{-1}$ | -0.09 | 0.14 | $5.27 \times 10^{-1}$ |
| Strelka:T4 | | | | | | | -0.08 | 0.10 | $3.95 \times 10^{-1}$ | 0.20 | 0.15 | $1.83 \times 10^{-1}$ | 0.16 | 0.14 | $2.55 \times 10^{-1}$ | 0.02 | 0.15 | $9.12 \times 10^{-1}$ | 0.04 | 0.14 | $7.75 \times 10^{-1}$ |
| Strelka:T5 | | | | | | | -0.37 | 0.10 | $8.92 \times 10^{-5}$ | -0.22 | 0.15 | $1.49 \times 10^{-1}$ | -0.30 | 0.15 | $3.72 \times 10^{-2}$ | -0.32 | 0.15 | $3.58 \times 10^{-2}$ | -0.31 | 0.15 | $3.21 \times 10^{-2}$ |
| Strelka:T6 | | | | | | | -0.15 | 0.09 | $1.14 \times 10^{-1}$ | 0.36 | 0.15 | $2.13 \times 10^{-2}$ | 0.33 | 0.15 | $2.44 \times 10^{-2}$ | -0.28 | 0.15 | $6.91 \times 10^{-2}$ | -0.22 | 0.15 | $1.32 \times 10^{-1}$ |
| Mutationseq:T3 | | | | | | | -0.18 | 0.10 | $5.60 \times 10^{-2}$ | -0.09 | 0.16 | $5.87 \times 10^{-1}$ | -0.07 | 0.15 | $6.17 \times 10^{-1}$ | -0.05 | 0.14 | $7.42 \times 10^{-1}$ | -0.09 | 0.14 | $5.15 \times 10^{-1}$ |
| Mutationseq:T4 | | | | | | | -0.17 | 0.10 | $8.14 \times 10^{-2}$ | -0.09 | 0.16 | $5.53 \times 10^{-1}$ | -0.06 | 0.15 | $6.75 \times 10^{-1}$ | -0.01 | 0.15 | $9.32 \times 10^{-1}$ | -0.04 | 0.14 | $7.53 \times 10^{-1}$ |
| Mutationseq:T5 | | | | | | | -0.44 | 0.10 | $4.01 \times 10^{-6}$ | -0.45 | 0.16 | $4.16 \times 10^{-3}$ | -0.47 | 0.15 | $1.58 \times 10^{-3}$ | -0.23 | 0.15 | $1.35 \times 10^{-1}$ | -0.28 | 0.14 | $5.10 \times 10^{-2}$ |
| Mutationseq:T6 | | | | | | | 0.03 | 0.09 | $7.72 \times 10^{-1}$ | -0.22 | 0.16 | $1.57 \times 10^{-1}$ | -0.50 | 0.15 | $8.18 \times 10^{-4}$ | -0.35 | 0.16 | $2.27 \times 10^{-2}$ | -0.35 | 0.15 | $1.52 \times 10^{-2}$ |
| phi | 22.37 | 1.76 | $6.54 \times 10^{-37}$ | | | | 33.03 | 2.18 | $5.48 \times 10^{-52}$ | 14.16 | 0.89 | $3.10 \times 10^{-57}$ | 16.14 | 1.02 | $6.81 \times 10^{-57}$ | 42.08 | 3.79 | $1.40 \times 10^{-28}$ | 46.35 | 4.14 | $4.36 \times 10^{-29}$ |
| Log-likelihood | 1506.53 | | | | | | 941.02 | | | 500.33 | | | 542.26 | | | 386.54 | | | 398.21 | | |
| Pseudo R-squared | 0.35 | | | | | | 0.68 | | | 0.83 | | | 0.85 | | | 0.81 | | | 0.83 | | |
