## Supplementary Table 3 for "Creating Standards for Evaluating Tumour Subclonal Reconstruction"

Supplementary Table 3: Alterative $\beta$ regressions for subchallenges 1C and 2A (N=500).

| $\beta$ | Estimate 1C | Std. Error 1C | P-value 1C | Estimate 2A | Std. Error 2A | P-value 2A |
|---|---|---|---|---|---|---|
| Intercept | 0.95 | 0.06 | $4.96 \times 10^{-61}$ | 0.12 | 0.09 | $1.92 \times 10^{-1}$ |
| Effective Depth | 0.09 | 0.02 | $2.73 \times 10^{-6}$ | 0.37 | 0.03 | $2.69 \times 10^{-32}$ |
| T3 | 0.00 | 0.07 | $9.93 \times 10^{-1}$ | -0.09 | 0.10 | $3.80 \times 10^{-1}$ |
| T4 | -0.12 | 0.07 | $7.12 \times 10^{-2}$ | 0.09 | 0.10 | $3.96 \times 10^{-1}$ |
| T5 | 0.06 | 0.07 | $3.84 \times 10^{-1}$ | 0.45 | 0.10 | $1.64 \times 10^{-5}$ |
| T6 | 0.04 | 0.07 | $5.76 \times 10^{-1}$ | 0.48 | 0.10 | $4.35 \times 10^{-6}$ |
| PhyloWGS | 0.07 | 0.02 | $4.74 \times 10^{-5}$ | 0.05 | 0.03 | $1.13 \times 10^{-1}$ |
| Downsampling | -0.02 | 0.02 | $3.09 \times 10^{-1}$ | -0.00 | 0.03 | $9.84 \times 10^{-1}$ |
| MuTect | 0.02 | 0.07 | $8.29 \times 10^{-1}$ | -0.33 | 0.11 | $3.14 \times 10^{-3}$ |
| SomaticSniper | 0.16 | 0.08 | $4.14 \times 10^{-2}$ | -0.54 | 0.12 | $1.25 \times 10^{-5}$ |
| Strelka | -0.02 | 0.07 | $7.50 \times 10^{-1}$ | -0.70 | 0.12 | $1.55 \times 10^{-9}$ |
| Mutationseq | 0.08 | 0.08 | $3.07 \times 10^{-1}$ | -0.92 | 0.12 | $1.17 \times 10^{-13}$ |
| Sensitivity | 0.23 | 0.02 | $4.26 \times 10^{-21}$ | 0.30 | 0.04 | $8.92 \times 10^{-13}$ |
| MuTect:T3 | -0.12 | 0.09 | $1.90 \times 10^{-1}$ | -0.04 | 0.14 | $7.58 \times 10^{-1}$ |
| MuTect:T4 | -0.24 | 0.09 | $8.05 \times 10^{-3}$ | -0.19 | 0.14 | $1.88 \times 10^{-1}$ |
| MuTect:T5 | -0.19 | 0.09 | $4.14 \times 10^{-2}$ | -0.18 | 0.15 | $2.07 \times 10^{-1}$ |
| MuTect:T6 | -0.07 | 0.09 | $4.73 \times 10^{-1}$ | 0.03 | 0.15 | $8.21 \times 10^{-1}$ |
| SomaticSniper:T3 | -0.35 | 0.09 | $1.15 \times 10^{-4}$ | -0.12 | 0.15 | $4.02 \times 10^{-1}$ |
| SomaticSniper:T4 | -0.17 | 0.09 | $7.12 \times 10^{-2}$ | -0.17 | 0.15 | $2.52 \times 10^{-1}$ |
| SomaticSniper:T5 | -0.51 | 0.09 | $2.56 \times 10^{-8}$ | -0.19 | 0.15 | $1.88 \times 10^{-1}$ |
| SomaticSniper:T6 | -0.10 | 0.09 | $3.11 \times 10^{-1}$ | 0.03 | 0.15 | $8.18 \times 10^{-1}$ |
| Strelka:T3 | -0.08 | 0.09 | $3.67 \times 10^{-1}$ | -0.02 | 0.15 | $8.97 \times 10^{-1}$ |
| Strelka:T4 | -0.20 | 0.09 | $3.24 \times 10^{-2}$ | 0.13 | 0.15 | $3.86 \times 10^{-1}$ |
| Strelka:T5 | -0.28 | 0.09 | $2.51 \times 10^{-3}$ | -0.07 | 0.15 | $6.46 \times 10^{-1}$ |
| Strelka:T6 | -0.24 | 0.09 | $8.63 \times 10^{-3}$ | 0.23 | 0.15 | $1.19 \times 10^{-1}$ |
| Mutationseq:T3 | -0.15 | 0.09 | $9.84 \times 10^{-2}$ | -0.09 | 0.15 | $5.56 \times 10^{-1}$ |
| Mutationseq:T4 | -0.19 | 0.09 | $3.95 \times 10^{-2}$ | -0.09 | 0.15 | $5.51 \times 10^{-1}$ |
| Mutationseq:T5 | -0.38 | 0.09 | $3.85 \times 10^{-5}$ | -0.37 | 0.15 | $1.38 \times 10^{-2}$ |
| Mutationseq:T6 | 0.03 | 0.09 | $7.75 \times 10^{-1}$ | -0.25 | 0.15 | $9.97 \times 10^{-2}$ |
| phi | 37.83 | 2.48 | $1.01 \times 10^{-52}$ | 15.97 | 1.01 | $8.62 \times 10^{-57}$ |
| Log-likelihood | 980.87 | | | 526.52 | | |
| Pseudo R-squared | 0.72 | | | 0.85 | | |
