## Supplementary Table 4 for "Creating Standards for Evaluating Tumour Subclonal Reconstruction"

Supplementary Table 4: Genaralized linear models for each subchallenge with the real tumour BAM for PD4120. $\beta$ regressions were used for all subchallenges except 1B where a binomial regression was used. N=100 for subchallenges 1 and 2 and N=50 for subchallenge 3.

| $\beta$ | Estimate 1A | Std. Error 1A | P-value 1A | Estimate 1B | Std. Error 1B | P-value 1B | Estimate 1C | Std. Error 1C | P-value 1C | Estimate 2A | Std. Error 2A | P-value 2A | Estimate 2B | Std. Error 2B | P-value 2B | Estimate 3A | Std. Error 3A | P-value 3A | Estimate 3B | Std. Error 3B | P-value 3B |
|---|---|---|---|---|---|---|---|---|---|---|---|---|---|---|---|---|---|---|---|---|---|
| Intercept | 1.97 | 0.09 | $7.31 \times 10^{-118}$ | 0.79 | 0.16 | $7.04 \times 10^{-7}$ | 0.87 | 0.05 | $1.60 \times 10^{-60}$ | 0.12 | 0.09 | $1.74 \times 10^{-1}$ | 0.44 | 0.10 | $1.65 \times 10^{-5}$ | 1.04 | 0.02 | $0.00 \times 10^{0}$ | 1.03 | 0.02 | $0.00 \times 10^{0}$ |
| Effective Depth | 0.09 | 0.03 | $4.20 \times 10^{-3}$ | 0.73 | 0.07 | $1.21 \times 10^{-27}$ | 0.15 | 0.02 | $1.91 \times 10^{-12}$ | 0.54 | 0.04 | $2.25 \times 10^{-42}$ | 0.66 | 0.04 | $9.07 \times 10^{-51}$ | 0.13 | 0.01 | $4.10 \times 10^{-46}$ | 0.12 | 0.01 | $1.08 \times 10^{-40}$ |
| PhyloWGS | -0.65 | 0.07 | $3.02 \times 10^{-23}$ | -0.57 | 0.13 | $9.87 \times 10^{-6}$ | -0.10 | 0.04 | $1.63 \times 10^{-2}$ | 0.29 | 0.08 | $1.16 \times 10^{-4}$ | 0.13 | 0.08 | $1.28 \times 10^{-1}$ | | | | | | |
| Downsampling | -0.00 | 0.06 | $9.85 \times 10^{-1}$ | 0.02 | 0.13 | $8.98 \times 10^{-1}$ | -0.01 | 0.04 | $8.77 \times 10^{-1}$ | -0.00 | 0.08 | $9.70 \times 10^{-1}$ | 0.02 | 0.08 | $8.36 \times 10^{-1}$ | 0.06 | 0.02 | $7.04 \times 10^{-4}$ | 0.05 | 0.02 | $4.20 \times 10^{-3}$ |
| SomaticSniper | -0.20 | 0.09 | $2.44 \times 10^{-2}$ | -0.39 | 0.18 | $3.11 \times 10^{-2}$ | -0.00 | 0.06 | $9.58 \times 10^{-1}$ | -0.33 | 0.11 | $1.99 \times 10^{-3}$ | -0.53 | 0.12 | $6.05 \times 10^{-6}$ | -0.12 | 0.02 | $2.89 \times 10^{-6}$ | -0.12 | 0.03 | $6.45 \times 10^{-6}$ |
| Strelka | -0.04 | 0.09 | $6.74 \times 10^{-1}$ | -0.07 | 0.18 | $7.16 \times 10^{-1}$ | -0.01 | 0.06 | $8.12 \times 10^{-1}$ | 0.10 | 0.11 | $3.36 \times 10^{-1}$ | 0.08 | 0.12 | $4.81 \times 10^{-1}$ | 0.02 | 0.03 | $3.64 \times 10^{-1}$ | 0.02 | 0.03 | $5.20 \times 10^{-1}$ |
| Mutationseq | -0.03 | 0.09 | $7.32 \times 10^{-1}$ | -0.10 | 0.18 | $5.85 \times 10^{-1}$ | 0.03 | 0.06 | $6.66 \times 10^{-1}$ | -0.12 | 0.11 | $2.75 \times 10^{-1}$ | -0.13 | 0.12 | $2.63 \times 10^{-1}$ | -0.07 | 0.03 | $3.51 \times 10^{-3}$ | -0.08 | 0.03 | $2.30 \times 10^{-3}$ |
| phi | 32.14 | 4.92 | $6.55 \times 10^{-11}$ | | | | 33.85 | 4.93 | $6.83 \times 10^{-12}$ | 11.47 | 1.61 | $9.88 \times 10^{-13}$ | 9.55 | 1.36 | $1.93 \times 10^{-12}$ | 523.47 | 107.02 | $1.00 \times 10^{-6}$ | 481.08 | 98.35 | $1.00 \times 10^{-6}$ |
| Log-likelihood | 197.56 | | | | | | 157.65 | | | 66.03 | | | 67.80 | | | 152.26 | | | 149.27 | | |
| Pseudo R-squared | 0.63 | | | | | | 0.33 | | | 0.69 | | | 0.72 | | | 0.84 | | | 0.81 | | |
