## Supplementary Table 5 for "Creating Standards for Evaluating Tumour Subclonal Reconstruction"

Supplementary Table 5: Genaralized linear models for each subchallenge with errors in the CNA-profiles. $\beta$ regressions were used for all subchallenges except 1B where a binomial regression was used. N=4250 for subchallenges 1 and 2 and N=2125 for subchallenge 3.

| $\beta$ | Estimate 1A | Std. Error 1A | P-value 1A | Estimate 1B | Std. Error 1B | P-value 1B | Estimate 1C | Std. Error 1C | P-value 1C | Estimate 2A | Std. Error 2A | P-value 2A | Estimate 2B | Std. Error 2B | P-value 2B | Estimate 3A | Std. Error 3A | P-value 3A | Estimate 3B | Std. Error 3B | P-value 3B |
|---|---|---|---|---|---|---|---|---|---|---|---|---|---|---|---|---|---|---|---|---|---|
| Intercept | 1.07 | 0.08 | $3.15 \times 10^{-42}$ | 0.70 | 0.19 | $1.73 \times 10^{-4}$ | 1.22 | 0.02 | $0.00 \times 10^{0}$ | 0.45 | 0.05 | $1.18 \times 10^{-18}$ | 0.34 | 0.05 | $7.35 \times 10^{-12}$ | 1.02 | 0.04 | $1.91 \times 10^{-164}$ | 1.01 | 0.03 | $2.47 \times 10^{-282}$ |
| Effective Depth | 0.03 | 0.01 | $2.63 \times 10^{-4}$ | 0.27 | 0.01 | $1.54 \times 10^{-94}$ | 0.25 | 0.01 | $5.55 \times 10^{-69}$ | 0.47 | 0.01 | $0.00 \times 10^{0}$ | 0.49 | 0.01 | $0.00 \times 10^{0}$ | 0.28 | 0.00 | $0.00 \times 10^{0}$ | 0.27 | 0.00 | $0.00 \times 10^{0}$ |
| T3 | -0.06 | 0.10 | $5.43 \times 10^{-1}$ | -0.04 | 0.06 | $5.19 \times 10^{-1}$ | -0.09 | 0.01 | $7.30 \times 10^{-16}$ | -0.05 | 0.02 | $1.94 \times 10^{-3}$ | 0.03 | 0.02 | $5.49 \times 10^{-2}$ | -0.04 | 0.05 | $3.84 \times 10^{-1}$ | -0.06 | 0.03 | $2.99 \times 10^{-2}$ |
| T4 | 0.01 | 0.10 | $8.92 \times 10^{-1}$ | -0.27 | 0.06 | $1.10 \times 10^{-5}$ | -0.27 | 0.01 | $1.24 \times 10^{-131}$ | 0.16 | 0.02 | $1.40 \times 10^{-21}$ | 0.11 | 0.02 | $1.26 \times 10^{-11}$ | -0.03 | 0.05 | $5.11 \times 10^{-1}$ | -0.05 | 0.03 | $1.13 \times 10^{-1}$ |
| T5 | -0.12 | 0.10 | $2.43 \times 10^{-1}$ | -0.45 | 0.06 | $6.79 \times 10^{-16}$ | -0.37 | 0.01 | $3.56 \times 10^{-261}$ | -0.12 | 0.02 | $3.85 \times 10^{-12}$ | -0.14 | 0.02 | $7.67 \times 10^{-18}$ | -0.11 | 0.05 | $3.27 \times 10^{-2}$ | -0.15 | 0.03 | $4.04 \times 10^{-7}$ |
| T6 | -0.24 | 0.11 | $2.73 \times 10^{-2}$ | -0.38 | 0.06 | $1.03 \times 10^{-10}$ | 0.08 | 0.01 | $5.11 \times 10^{-14}$ | 0.57 | 0.02 | $1.55 \times 10^{-262}$ | 0.64 | 0.02 | $0.00 \times 10^{0}$ | 0.15 | 0.05 | $2.83 \times 10^{-3}$ | 0.21 | 0.03 | $1.35 \times 10^{-12}$ |
| PhyloWGS | 0.19 | 0.02 | $7.80 \times 10^{-28}$ | 0.05 | 0.04 | $2.25 \times 10^{-1}$ | 0.04 | 0.01 | $6.06 \times 10^{-10}$ | 0.07 | 0.01 | $1.85 \times 10^{-10}$ | 0.13 | 0.01 | $2.00 \times 10^{-35}$ | | | | | | |
| MuTect | -0.01 | 0.03 | $8.21 \times 10^{-1}$ | -0.60 | 0.23 | $9.92 \times 10^{-3}$ | -0.32 | 0.01 | $1.82 \times 10^{-188}$ | -0.71 | 0.07 | $6.31 \times 10^{-26}$ | -0.70 | 0.07 | $1.56 \times 10^{-26}$ | -0.43 | 0.01 | $1.30 \times 10^{-275}$ | -0.43 | 0.03 | $1.19 \times 10^{-48}$ |
| SomaticSniper | -0.01 | 0.03 | $7.09 \times 10^{-1}$ | -0.15 | 0.30 | $6.10 \times 10^{-1}$ | -0.32 | 0.01 | $2.60 \times 10^{-190}$ | -1.09 | 0.07 | $3.34 \times 10^{-57}$ | -1.09 | 0.07 | $8.98 \times 10^{-61}$ | -0.68 | 0.01 | $0.00 \times 10^{0}$ | -0.70 | 0.03 | $4.16 \times 10^{-122}$ |
| Strelka | -0.04 | 0.03 | $1.78 \times 10^{-1}$ | -0.33 | 0.24 | $1.62 \times 10^{-1}$ | -0.39 | 0.01 | $1.69 \times 10^{-276}$ | -1.02 | 0.07 | $4.48 \times 10^{-51}$ | -0.92 | 0.07 | $4.62 \times 10^{-44}$ | -0.61 | 0.01 | $0.00 \times 10^{0}$ | -0.64 | 0.03 | $2.47 \times 10^{-104}$ |
| Mutationseq | 0.01 | 0.03 | $7.82 \times 10^{-1}$ | -0.67 | 0.23 | $3.90 \times 10^{-3}$ | -0.40 | 0.01 | $1.32 \times 10^{-296}$ | -1.59 | 0.07 | $5.03 \times 10^{-114}$ | -1.54 | 0.07 | $1.16 \times 10^{-112}$ | -0.83 | 0.01 | $0.00 \times 10^{0}$ | -0.78 | 0.03 | $5.90 \times 10^{-149}$ |
| ploidy x2 | -0.50 | 0.11 | $1.26 \times 10^{-5}$ | -0.49 | 0.24 | $4.09 \times 10^{-2}$ | -0.20 | 0.02 | $1.04 \times 10^{-22}$ | -0.30 | 0.07 | $1.12 \times 10^{-5}$ | -0.42 | 0.07 | $2.63 \times 10^{-10}$ | 0.29 | 0.05 | $5.87 \times 10^{-9}$ | 0.08 | 0.02 | $1.02 \times 10^{-3}$ |
| scramble | -0.30 | 0.08 | $3.79 \times 10^{-4}$ | -0.23 | 0.20 | $2.42 \times 10^{-1}$ | 0.01 | 0.02 | $4.17 \times 10^{-1}$ | -0.28 | 0.05 | $2.82 \times 10^{-7}$ | -0.24 | 0.05 | $5.68 \times 10^{-6}$ | -0.02 | 0.04 | $5.84 \times 10^{-1}$ | -0.07 | 0.02 | $4.51 \times 10^{-4}$ |
| scramble gains | -0.26 | 0.08 | $1.89 \times 10^{-3}$ | -0.19 | 0.20 | $3.24 \times 10^{-1}$ | -0.02 | 0.02 | $2.72 \times 10^{-1}$ | -0.30 | 0.05 | $2.37 \times 10^{-8}$ | -0.29 | 0.05 | $5.28 \times 10^{-8}$ | -0.03 | 0.04 | $4.63 \times 10^{-1}$ | -0.04 | 0.02 | $4.14 \times 10^{-2}$ |
| scramble loss | 0.29 | 0.08 | $4.80 \times 10^{-4}$ | 0.02 | 0.20 | $9.17 \times 10^{-1}$ | 0.04 | 0.02 | $2.11 \times 10^{-2}$ | -0.11 | 0.05 | $3.53 \times 10^{-2}$ | -0.07 | 0.05 | $2.00 \times 10^{-1}$ | -0.09 | 0.04 | $2.25 \times 10^{-2}$ | -0.10 | 0.02 | $5.76 \times 10^{-7}$ |
| proportion CNAs scrambled | -0.15 | 0.01 | $1.75 \times 10^{-56}$ | -0.00 | 0.00 | $5.19 \times 10^{-1}$ | -0.03 | 0.00 | $1.36 \times 10^{-12}$ | -0.03 | 0.01 | $5.16 \times 10^{-6}$ | -0.03 | 0.01 | $3.60 \times 10^{-6}$ | 0.02 | 0.00 | $4.49 \times 10^{-6}$ | 0.03 | 0.00 | $1.34 \times 10^{-11}$ |
| T3:ploidy x2 | 0.04 | 0.16 | $8.10 \times 10^{-1}$ | | | | | | | | | | | | | -0.31 | 0.07 | $9.13 \times 10^{-6}$ | | | |
| T3:scramble | 0.16 | 0.12 | $1.81 \times 10^{-1}$ | | | | | | | | | | | | | -0.13 | 0.05 | $1.43 \times 10^{-2}$ | | | |
| T3:scramble gains | 0.14 | 0.12 | $2.45 \times 10^{-1}$ | | | | | | | | | | | | | 0.05 | 0.05 | $3.71 \times 10^{-1}$ | | | |
| T3:scramble loss | -0.00 | 0.11 | $9.95 \times 10^{-1}$ | | | | | | | | | | | | | -0.04 | 0.05 | $4.30 \times 10^{-1}$ | | | |
| T4:ploidy x2 | 0.24 | 0.16 | $1.34 \times 10^{-1}$ | | | | | | | | | | | | | -0.25 | 0.07 | $2.75 \times 10^{-4}$ | | | |
| T4:scramble | 0.04 | 0.12 | $7.33 \times 10^{-1}$ | | | | | | | | | | | | | 0.04 | 0.05 | $5.12 \times 10^{-1}$ | | | |
| T4:scramble gains | 0.08 | 0.11 | $4.77 \times 10^{-1}$ | | | | | | | | | | | | | 0.04 | 0.05 | $5.16 \times 10^{-1}$ | | | |
| T4:scramble loss | 0.03 | 0.11 | $7.89 \times 10^{-1}$ | | | | | | | | | | | | | 0.07 | 0.05 | $2.24 \times 10^{-1}$ | | | |
| T5:ploidy x2 | 0.18 | 0.16 | $2.62 \times 10^{-1}$ | | | | | | | | | | | | | -0.18 | 0.07 | $1.06 \times 10^{-2}$ | | | |
| T5:scramble | 0.52 | 0.12 | $8.27 \times 10^{-6}$ | | | | | | | | | | | | | 0.03 | 0.05 | $5.61 \times 10^{-1}$ | | | |
| T5:scramble gains | 0.42 | 0.12 | $2.69 \times 10^{-4}$ | | | | | | | | | | | | | 0.07 | 0.05 | $1.83 \times 10^{-1}$ | | | |
| T5:scramble loss | 0.04 | 0.11 | $7.11 \times 10^{-1}$ | | | | | | | | | | | | | 0.10 | 0.05 | $6.93 \times 10^{-2}$ | | | |
| T6:ploidy x2 | 0.30 | 0.16 | $6.40 \times 10^{-2}$ | | | | | | | | | | | | | -0.35 | 0.07 | $4.28 \times 10^{-7}$ | | | |
| T6:scramble | 0.52 | 0.12 | $1.01 \times 10^{-5}$ | | | | | | | | | | | | | -0.02 | 0.05 | $7.24 \times 10^{-1}$ | | | |
| T6:scramble gains | 0.33 | 0.12 | $5.19 \times 10^{-3}$ | | | | | | | | | | | | | -0.03 | 0.05 | $5.99 \times 10^{-1}$ | | | |
| T6:scramble loss | 0.05 | 0.12 | $6.41 \times 10^{-1}$ | | | | | | | | | | | | | 0.07 | 0.05 | $1.94 \times 10^{-1}$ | | | |
| MuTect:ploidy x2 | | | | 0.10 | 0.31 | $7.62 \times 10^{-1}$ | | | | 0.15 | 0.09 | $1.23 \times 10^{-1}$ | 0.23 | 0.09 | $1.44 \times 10^{-2}$ | | | | | | |
| MuTect:scramble | | | | 0.24 | 0.25 | $3.38 \times 10^{-1}$ | | | | 0.18 | 0.07 | $1.23 \times 10^{-2}$ | 0.15 | 0.07 | $3.81 \times 10^{-2}$ | | | | | | |
| MuTect:scramble gains | | | | 0.18 | 0.25 | $4.87 \times 10^{-1}$ | | | | 0.21 | 0.07 | $4.35 \times 10^{-3}$ | 0.19 | 0.07 | $8.26 \times 10^{-3}$ | | | | | | |
| MuTect:scramble loss | | | | 0.02 | 0.26 | $9.53 \times 10^{-1}$ | | | | 0.12 | 0.07 | $1.13 \times 10^{-1}$ | 0.08 | 0.07 | $2.41 \times 10^{-1}$ | | | | | | |
| SomaticSniper:ploidy x2 | | | | -0.41 | 0.39 | $2.87 \times 10^{-1}$ | | | | 0.25 | 0.10 | $9.48 \times 10^{-3}$ | 0.35 | 0.09 | $1.61 \times 10^{-4}$ | | | | | | |
| SomaticSniper:scramble | | | | 0.31 | 0.32 | $3.33 \times 10^{-1}$ | | | | 0.26 | 0.07 | $4.91 \times 10^{-4}$ | 0.25 | 0.07 | $4.97 \times 10^{-4}$ | | | | | | |
| SomaticSniper:scramble gains | | | | 0.14 | 0.32 | $6.63 \times 10^{-1}$ | | | | 0.30 | 0.07 | $6.12 \times 10^{-5}$ | 0.30 | 0.07 | $2.90 \times 10^{-5}$ | | | | | | |
| SomaticSniper:scramble loss | | | | 0.14 | 0.32 | $6.56 \times 10^{-1}$ | | | | 0.13 | 0.07 | $9.06 \times 10^{-2}$ | 0.10 | 0.07 | $1.52 \times 10^{-1}$ | | | | | | |
| Strelka:ploidy x2 | | | | 0.36 | 0.32 | $2.69 \times 10^{-1}$ | | | | 0.44 | 0.09 | $3.28 \times 10^{-6}$ | 0.42 | 0.09 | $5.83 \times 10^{-6}$ | | | | | | |
| Strelka:scramble | | | | 0.22 | 0.26 | $4.07 \times 10^{-1}$ | | | | 0.34 | 0.07 | $3.80 \times 10^{-6}$ | 0.27 | 0.07 | $1.98 \times 10^{-4}$ | | | | | | |
| Strelka:scramble gains | | | | 0.16 | 0.26 | $5.49 \times 10^{-1}$ | | | | 0.41 | 0.07 | $3.90 \times 10^{-8}$ | 0.32 | 0.07 | $1.00 \times 10^{-5}$ | | | | | | |
| Strelka:scramble loss | | | | -0.08 | 0.26 | $7.66 \times 10^{-1}$ | | | | 0.19 | 0.07 | $1.04 \times 10^{-2}$ | 0.11 | 0.07 | $1.13 \times 10^{-1}$ | | | | | | |
| Mutationseq:ploidy x2 | | | | -0.02 | 0.31 | $9.51 \times 10^{-1}$ | | | | 0.45 | 0.10 | $4.25 \times 10^{-6}$ | 0.51 | 0.10 | $6.77 \times 10^{-8}$ | | | | | | |
| Mutationseq:scramble | | | | 0.35 | 0.25 | $1.64 \times 10^{-1}$ | | | | 0.32 | 0.08 | $3.00 \times 10^{-5}$ | 0.30 | 0.07 | $6.37 \times 10^{-5}$ | | | | | | |
| Mutationseq:scramble gains | | | | 0.27 | 0.25 | $2.81 \times 10^{-1}$ | | | | 0.37 | 0.08 | $1.39 \times 10^{-6}$ | 0.36 | 0.07 | $1.96 \times 10^{-6}$ | | | | | | |
| Mutationseq:scramble loss | | | | 0.07 | 0.25 | $7.84 \times 10^{-1}$ | | | | 0.15 | 0.08 | $5.71 \times 10^{-2}$ | 0.12 | 0.07 | $1.18 \times 10^{-1}$ | | | | | | |
| MuTect:T3 | | | | | | | | | | | | | | | | | | | 0.00 | 0.04 | $9.87 \times 10^{-1}$ |
| MuTect:T4 | | | | | | | | | | | | | | | | | | | -0.08 | 0.04 | $5.16 \times 10^{-2}$ |
| MuTect:T5 | | | | | | | | | | | | | | | | | | | -0.09 | 0.04 | $2.61 \times 10^{-2}$ |
| MuTect:T6 | | | | | | | | | | | | | | | | | | | -0.04 | 0.04 | $3.46 \times 10^{-1}$ |
| SomaticSniper:T3 | | | | | | | | | | | | | | | | | | | -0.09 | 0.04 | $3.83 \times 10^{-2}$ |
| SomaticSniper:T4 | | | | | | | | | | | | | | | | | | | 0.04 | 0.04 | $2.78 \times 10^{-1}$ |
| SomaticSniper:T5 | | | | | | | | | | | | | | | | | | | 0.02 | 0.04 | $7.15 \times 10^{-1}$ |
| SomaticSniper:T6 | | | | | | | | | | | | | | | | | | | -0.10 | 0.04 | $2.08 \times 10^{-2}$ |
| Strelka:T3 | | | | | | | | | | | | | | | | | | | -0.04 | 0.04 | $3.92 \times 10^{-1}$ |
| Strelka:T4 | | | | | | | | | | | | | | | | | | | 0.08 | 0.04 | $6.60 \times 10^{-2}$ |
| Strelka:T5 | | | | | | | | | | | | | | | | | | | -0.04 | 0.04 | $3.54 \times 10^{-1}$ |
| Strelka:T6 | | | | | | | | | | | | | | | | | | | -0.01 | 0.04 | $7.71 \times 10^{-1}$ |
| Mutationseq:T3 | | | | | | | | | | | | | | | | | | | -0.07 | 0.04 | $1.17 \times 10^{-1}$ |
| Mutationseq:T4 | | | | | | | | | | | | | | | | | | | 0.01 | 0.04 | $8.79 \times 10^{-1}$ |
| Mutationseq:T5 | | | | | | | | | | | | | | | | | | | -0.00 | 0.04 | $9.95 \times 10^{-1}$ |
| Mutationseq:T6 | | | | | | | | | | | | | | | | | | | -0.17 | 0.04 | $3.51 \times 10^{-5}$ |
| Effective Depth:ploidy x2 | | | | | | | -0.17 | 0.02 | $4.06 \times 10^{-17}$ | | | | | | | | | | | | |
| Effective Depth:scramble | | | | | | | -0.01 | 0.02 | $3.69 \times 10^{-1}$ | | | | | | | | | | | | |
| Effective Depth:scramble gains | | | | | | | -0.02 | 0.02 | $2.16 \times 10^{-1}$ | | | | | | | | | | | | |
| Effective Depth:scramble loss | | | | | | | 0.00 | 0.02 | $7.64 \times 10^{-1}$ | | | | | | | | | | | | |
| phi | 2.80 | 0.07 | $0.00 \times 10^{0}$ | | | | 29.78 | 0.67 | $0.00 \times 10^{0}$ | 14.59 | 0.31 | $0.00 \times 10^{0}$ | 15.50 | 0.34 | $0.00 \times 10^{0}$ | 54.84 | 1.74 | $2.76 \times 10^{-219}$ | 46.82 | 1.46 | $1.05 \times 10^{-226}$ |
| Log-likelihood | 7042.39 | | | | | | 7081.81 | | | 3939.12 | | | 4132.92 | | | 3380.31 | | | 3224.20 | | |
| Pseudo R-squared | 0.31 | | | | | | 0.61 | | | 0.79 | | | 0.80 | | | 0.84 | | | 0.83 | | |
